# CD5L constraints acute and systemic inflammation and can be a novel potent therapeutic agent against sepsis

**DOI:** 10.1101/2022.03.08.483540

**Authors:** Liliana Oliveira, Ana P. Gomes, Rita F. Santos, Marcos S. Cardoso, Ana Nóvoa, Hervé Luche, Fátima Gartner, Bernard Malissen, Moisés Mallo, Alexandre M. Carmo

## Abstract

The global burden of sepsis, with an estimated 49 million cases and 11 million deaths in 2017, often passes unnoticed to the general public even though it is the direct cause of nearly 20% of all deaths worldwide. This unawareness is perhaps due to misconceptions, or miscoding in the reporting of the ultimate causes of death, as in many diseases it is not the actual infectious agent that causes the biggest harm. Rather, it is the uncontrolled inflammation leading to septic shock that is the most menacing manifestation associated with many infections, and becomes deadly serious once it has passed the stage where anti-microbial drugs no longer have any effect to inactivate or destroy the pathogen. Here we show that the combined anti-bacterial and anti-inflammatory properties of the scavenger receptor cysteine-rich (SRCR) protein CD5L contribute to a remarkable therapeutic effect of the protein to fight sepsis, such that when exogenously administered in C57BL/6 mice with induced lethal-grade sepsis, it can be a very effective curative agent to treat this condition. The resistance conferred by CD5L to polybacterial-induced sepsis using the cecal ligation and puncture (CLP) model is consistent with the reported observations that CD5L physically binds and inactivates diverse species and strains of bacteria. Accordingly, our CD5L-knockout mice are significantly more susceptible to experimentally-induced mid-grade CLP than wild-type animals. We show that CD5L is centered on promoting neutrophil recruitment and activation, overall contributing to reducing the bacteria burden of the animals. However, the dramatic susceptibility of CD5L-deficient animals is not necessarily correlated only with pathogen load, as these mice are also extremely susceptible to sterile sepsis induced by nonlethal doses of LPS. Notwithstanding the observed capacity of CD5L to directly bind to a broad range of pathogens, typical of many PRRs, our evidence suggests that the anti-inflammatory properties of the protein are at least as important as its pathogen-binding potential, and can, and should, be explored to treat the deadly inflammation storm that is sepsis.

## Introduction

SRCR proteins are a conserved family of extracellular or membrane-bound receptors present in all metazoans. Individual SRCR proteins display varied roles in, *e.g*., cell differentiation, homeostasis or apoptosis, but only now some key features are emerging as distinctive of this family. CD6 and CD163, receptors of T cells and macrophages, respectively, DMBT1, broadly expressed, and CD5L, SSC4D, SSC5D, which are blood circulating proteins, all are molecular sensors of Grampositive and Gram-negative bacteria [1–3]. Given that some of the phylogenetically more ancient SRCR proteins like MARCO, SR-AI and SCARA-5, contribute to bacteria recognition and clearance [1], the concept is arising that SRCRs constitute an autonomous family of pattern recognition receptors (PRR). However, unlike other groups of PRRs such as toll-like receptors and nucleotide-binding oligomerization domain-like receptors, SRCR proteins may actively contribute to the control and resolution of inflammation [4]. For example, SR-AI efficiently opsonizes *N. meningitidis* without promoting significant inflammatory cytokine secretion [5], CD5 and CD6 refrain the strength of T cell responses [6, 7], CD163, a marker of M2 macrophages, induces the expression of anti-inflammatory IL-10 [8]; and the absence of SCARA-5 results in autoimmune disease [9].

CD5 antigen-like (CD5L), also known as Spa or apoptosis inhibitor of macrophages (AIM), is produced by tissue macrophages [10] and is being acknowledge as a multifunctional protein involved in different processes such as metabolism, infection and autophagy [11]. A common denominator in all these contexts is the capacity of CD5L to establish interactions with multiple surfaces, namely of pathogens, cellular debris, and different synthetic particles, all to be phagocyted [12–16]. By helping to remove dead cells and damage-associated molecular patterns (DAMPs), CD5L also contributes to decreasing the sterile inflammation associated with these forms and thus serves as a facilitator of tissue repair and healing in different *in vivo* models [14, 17–19].

Notwithstanding some degree of controversy, CD5L is also being associated with the downmodulation of inflammatory responses in infection models, either by improving the microbicidal capacity of macrophages infected by, *e.g*., *L. monocytogenes* [20] or *M. tuberculosis* [21], or acting directly on the mechanisms of inflammation by reducing macrophage secretion of TNF- α, IL-1β and IL-6 [12, 16], restraining the pro-inflammatory signature in non-pathogenic Th17 cells [22], and ameliorating fungus-induced peritoneal injury [19]. In a recent study of a mouse model of sepsis induced by CLP, the upregulation of CD5L gene expression, resulting from the administration of recombinant growth/differentiation factor 3, was suggested to be the key factor in improving mouse survival, along with significant reductions in bacterial load, pro-inflammatory cytokine levels and organ damage [23]. However, these results are at odds with a previous report describing that CD5L administration at the time of CLP induction aggravated the inflammation and pathology in septic mice, and overall increased the mortality [24]. In the present study we attempted to address the divergent aspects of the role of CD5L by evaluating on one hand the susceptibility of CD5L-deficient mice to CLP-induced mid-grade sepsis, and on the other assess the therapeutic value of exogenously administered recombinant CD5L (rCD5L) to fight full overt sepsis induced by lethal-grade CLP on C57BL/6 WT mice.

According to The Third International Consensus Definitions Task Force, sepsis is defined as “a life-threatening organ dysfunction caused by a dysregulated host response to infection”, and septic shock is “a subset of sepsis in which particularly profound circulatory, cellular, and metabolic abnormalities are associated with a greater risk of mortality than with sepsis alone” [25]. Usually, causative microbial foci trigger a disproportionate production of pro-inflammatory mediators resulting in a massive inflammatory response. However, the actual infection may not constitute the biggest risk; the uncontrolled inflammation leading to septic shock is the most threatening indicator of the syndrome and the effectiveness of antibiotics to treat this condition may be limited. The described anti-microbial and anti-inflammatory properties of CD5L suit our purpose to test its therapeutic application against sepsis, using the gold standard model of sepsis in mice. By providing a therapeutic scheme that intently reproduces the treatments applied in the clinic, we here provide relevant data showing the potential of CD5L as a powerful new agent to treat sepsis of bacterial origin.

## Results

### CD5L-deficient mice are susceptible and have decreased survival to non-lethal forms of sepsis

CD5L-knockout (CD5L-KO) mice were generated by CRISPR/Cas9 engineering through targeting the SRCR domain 1-encoding exon 3 of the *Cd5l* gene in C57BL/6 mice, with the insertion of three in-frame stop codons and a frame shift (**Fig. 1A**). Mice were viable and healthy, and at 12 weeks of age an analysis of the number of leukocytes, frequency of sub-populations and several hematological parameters, obtained through comprehensive immunophenotyping of the spleen, thymus, peripheral blood and peritoneal cavity, showed no relevant differences between WT and CD5L-KO mice in any of the parameters evaluated in all organs tested (**Fig. S1**).

**Figure 1.**
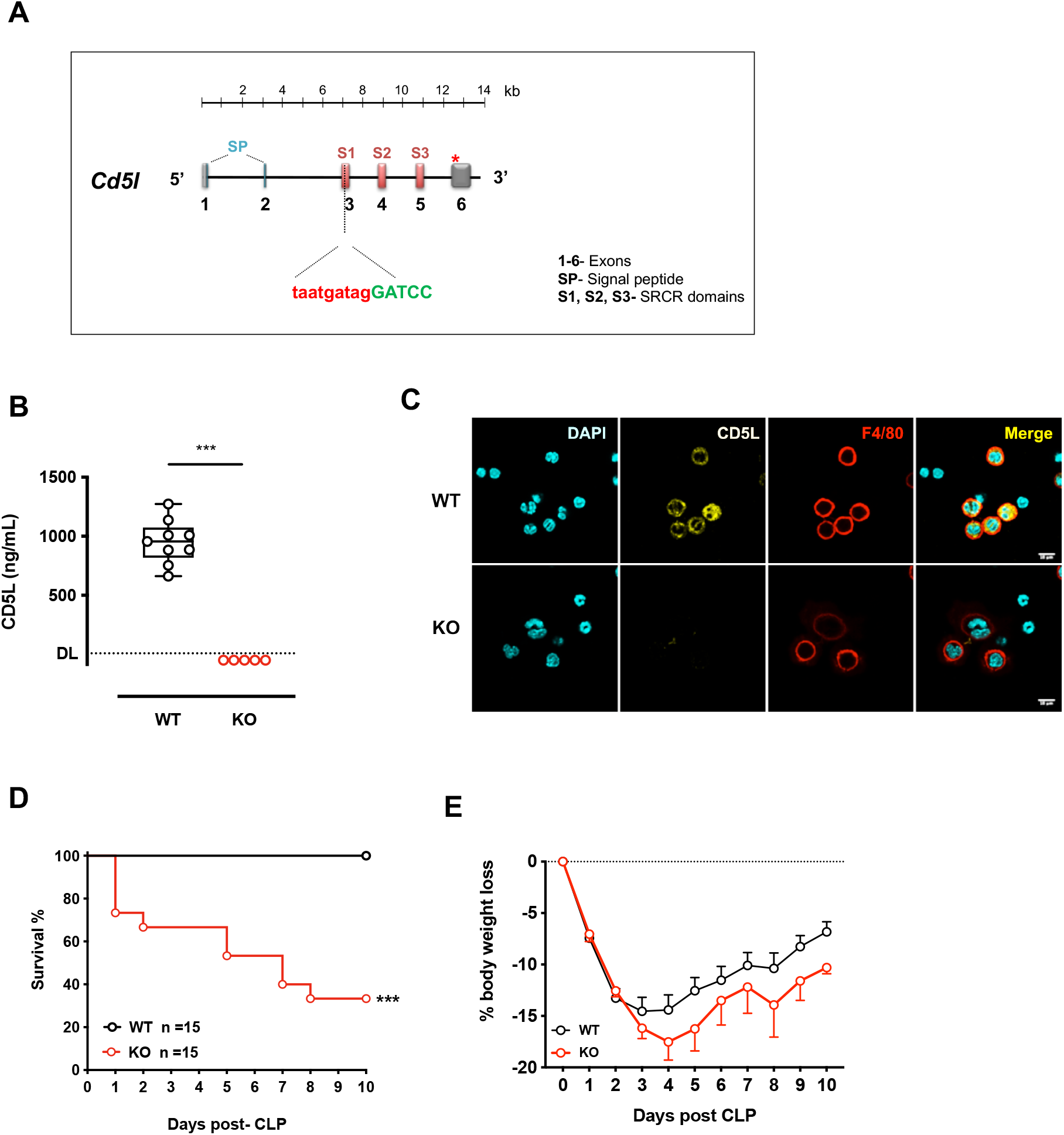
CD5L-KO mouse are susceptible to non-lethal forms of sepsis induced by LPS or CLP. **(A)** CD5L-KO mice were generated by CRISPR-Cas9 engineering. Schematic representation of the genome editing strategy to silence *Cd5l* expression by inserting three in-frame stop codons and a frame shift. **(B)** Quantification by ELISA of CD5L in the sera of C57BL/6 WT and CD5L-KO mice. DL: detection limit (6.25 pg/mL). Data from at least 3 independent experiments (Mann-Whitney test). **(C)** Intracellular CD5L was detected, after permeabilization of the cells with Triton X100, by staining with goat anti-CD5L polyclonal antibody followed by Alexa 488-coupled donkey anti-goat secondary antibody. Peritoneal macrophages from WT or CD5L-KO healthy mice were identified through staining with anti-mouse F4/80 mAb followed by Alexa 594-coupled anti-rat secondary antibody. DAPI was used as a nuclear counterstaining. Scale bar: 10 μm. **(D)** WT or CD5L-KO mice were subjected to surgery to expose and ligate the cecum at approximately half the distance between the distal pole and the base of the cecum, and through-and-through puncture with a 21G needle, to induce a mid-grade sepsis condition. N = 15 (each group). Kaplan–Meier survival curves were generated to compare mortality between the two groups and significance was determined by log-rank (Mantel-Cox) test. **(E)** Body weight loss quantification. ***, *p* < 0.005.

Circulating CD5L was present at high levels in WT mice, as measured by ELISA, but was not detectable in the blood of CD5L-deficient animals (**Fig. 1B**). The main source of CD5L is believed to be tissue macrophages [10], and whereas peritoneal F4/80^+^ macrophages of WT mice expressed high levels of the protein, as detected by immunofluorescence, equivalent cells from CD5L-KO mice lacked the expression of CD5L (**Fig. 1C**).

Notably, CD5L deficiency resulted in a higher susceptibility to a mid-grade level of polymicrobial infection through a cecal ligation and puncture (CLP) surgical procedure, and while 100% of WT mice survived the process, more than 60% of CD5L-KO mice succumbed after developing sepsis (**Fig. 1D**). Furthermore, CD5L-KO mice showed greater weight loss overtime comparing with WT mice, indicating a worse general condition (**Fig. 1E**).

### CD5L-deficient mice have an impaired immune response to sepsis

The infection that followed CLP and the elicited immune response were characterized at 6, 24 and 72 h after the surgery in WT and CD5L-KO mice. Counts of colony-forming units (CFU) showed that CD5L-KO mice presented faster increases in bacteria levels, compared with WT mice, and after 72 h of CLP displayed significant blood bacteremia, contrasting with the effective control of the systemic infection by WT mice (**Fig. 2A**). In line with this observation, CD5L-KO mice showed significantly increased bacterial counts in the three organs analyzed, lungs, liver and kidneys at 72 h post-surgery, indicating an impairment in bacterial control.

**Figure 2.**
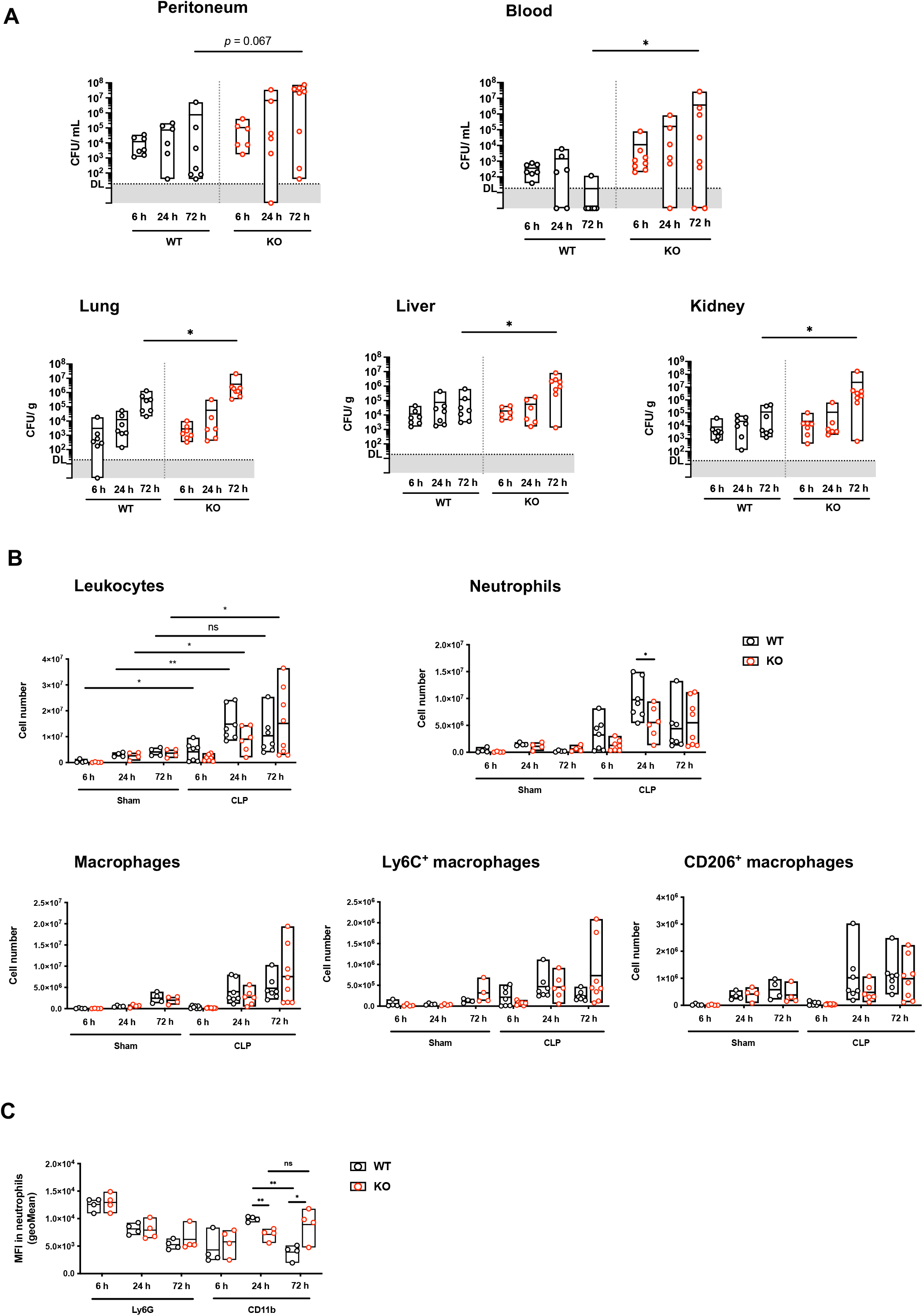
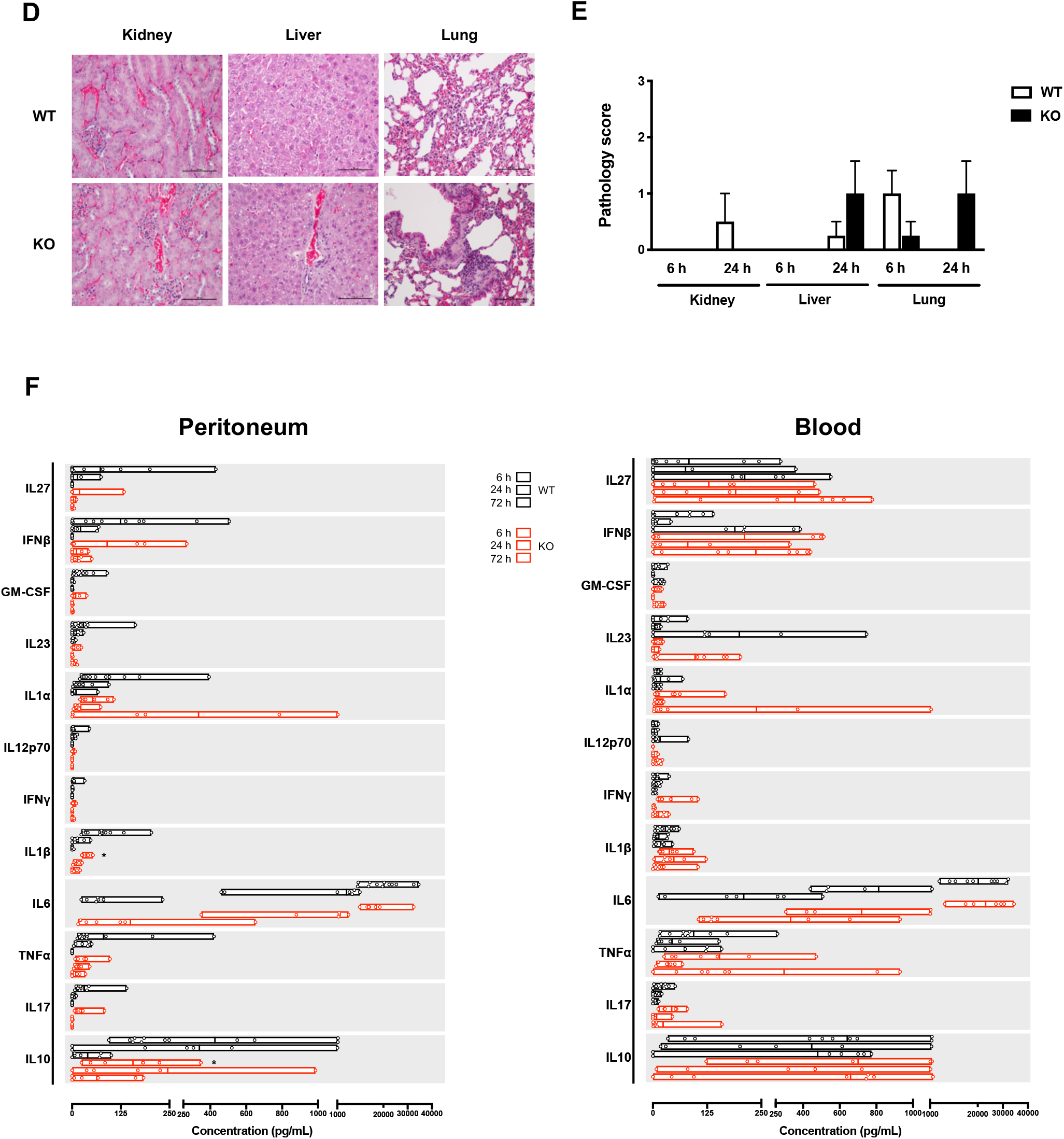
CD5L-KO mouse have an impaired immune response to polymicrobial-induced sepsis. WT or CD5L-KO mice were subjected to CLP to induce mid-grade sepsis. For that, animals were subjected to surgery, exposure and ligation of the cecum (corresponding to approximately half the distance between the distal pole and the base of the cecum) and through-and-through puncture with a 21G needle. Control mice (sham) were subjected to the same surgical procedure but without cecum ligation and puncture. Mice were euthanized 6 or 24 h post-surgery. **(A)** CFU counts from the peritoneal cavity, blood, lung, liver and kidney. **(B)** Absolute counts assessed by flow cytometry of cellular populations in the peritoneal cavity. Subsets analyzed: leukocytes, CD45^+^; neutrophils, CD45^+^CD11b^+^Ly6G^+^; macrophages, CD45^+^CD11b^+^CD11c^-^F4/80^+^. Ly6C and CD206 markers were used to distinguish between M1 and M2 within the macrophage population. **(C)** Ly6G and CD11b mean fluorescence intensity (MFI) in expressing neutrophils obtained by geometric mean statistics. **(D)** Lung, liver and kidney tissue sections obtained 24 h post CLP were stained with hematoxylin and eosin. Scale: 100 μm **(E)** Pathology score of lung, liver and kidney 24 h post CLP in a double blinded analysis. 0, no signs of inflammation; 1, minimal inflammatory signs; 2, mild inflammation; 3, moderate to severe inflammation. **(F)** The indicated cytokines were quantified by a multiplex bead-based immunoassay in samples from the peritoneal cavity (left panel) and blood serum (right panel). Data from at least 2 independent experiments (Mann-Whitney test). *, *p* < 0.05; **, *p* < 0.01; ns, not statistically significant.

On CLP, there was a rapid recruitment of leukocytes to the peritoneal cavity but CD5L-KO mice had already at 6 h a visibly smaller number of infiltrating cells than WT mice, which was significantly more pronounced at the 24 h time point. This difference was mostly attributable to a reduced number (**Fig. 2B**), though not of percentage (**Fig. S2**), of neutrophils being recruited into the peritoneal cavity 24 h after CLP. Importantly, in addition to the impaired recruitment, neutrophils of the CD5L-KO mice displayed, compared with WT mice, a slow growing but reduced surface expression of the α-integrin CD11b, consistent with a less activated state, while the expression of Ly6G remained unchanged between WT and CD5L-KO mice (**Fig. 2C**). By contrast, WT neutrophils showed a fast increase of CD11b expression at 24 h, and then an also rapid decline of the marker, consistent with a resolution phase of the inflammatory response.

CD5L has been described to be involved in the acquisition of the M2 phenotype by macrophages [16]. We therefore investigated whether there was an imbalance between M1 and M2 within the peritoneal macrophage population, using the classical markers Ly6C and CD206, respectively. However, no major differences in the M1 and M2 infiltrating populations were observed between WT and CD5L-KO mice (**Fig. 2B**).

At 24 h post-CLP, CD5L-KO mice showed increased signs of pathology in the lungs and liver, comparing with WT mice (**Fig. 2D, 2E**). A multiplex analysis of important inflammation-related cytokines in the peritoneal fluids and blood revealed that CD5L-KO mice produced significantly lower amounts of IL-1β and IL-10 than WT mice, but only in the peritoneal cavity and at 6 h after CLP **(Fig. 2F)**.

Overall, our results showed that CD5L-KO mice have a striking defect in fighting sepsis and display insufficiencies in the recruitment and activation of neutrophils. We thus needed to assess whether the impaired immune response could be due either to deficiencies in function or to the deficit in the number of recruited neutrophils, or both, and what could be the role of CD5L in these processes.

### CD5L levels change upon infection

Upon CLP aggression, the natural levels of CD5L increased in the peritoneal cavity of WT mice, peaking at 6 h after CLP and then receding at 24 h (**Fig. 3A, left panel**). By contrast, the levels of CD5L in the blood showed the opposite behavior, declining at 6 h and returning to normal after 24 h (**Fig. 3A, right panel**). We hypothesized that those two phenomena could be linked and that the decrease in the circulating levels of CD5L was the result of the redeployment of the protein from circulation, namely the mobilization to the site of aggression.

**Figure 3.**
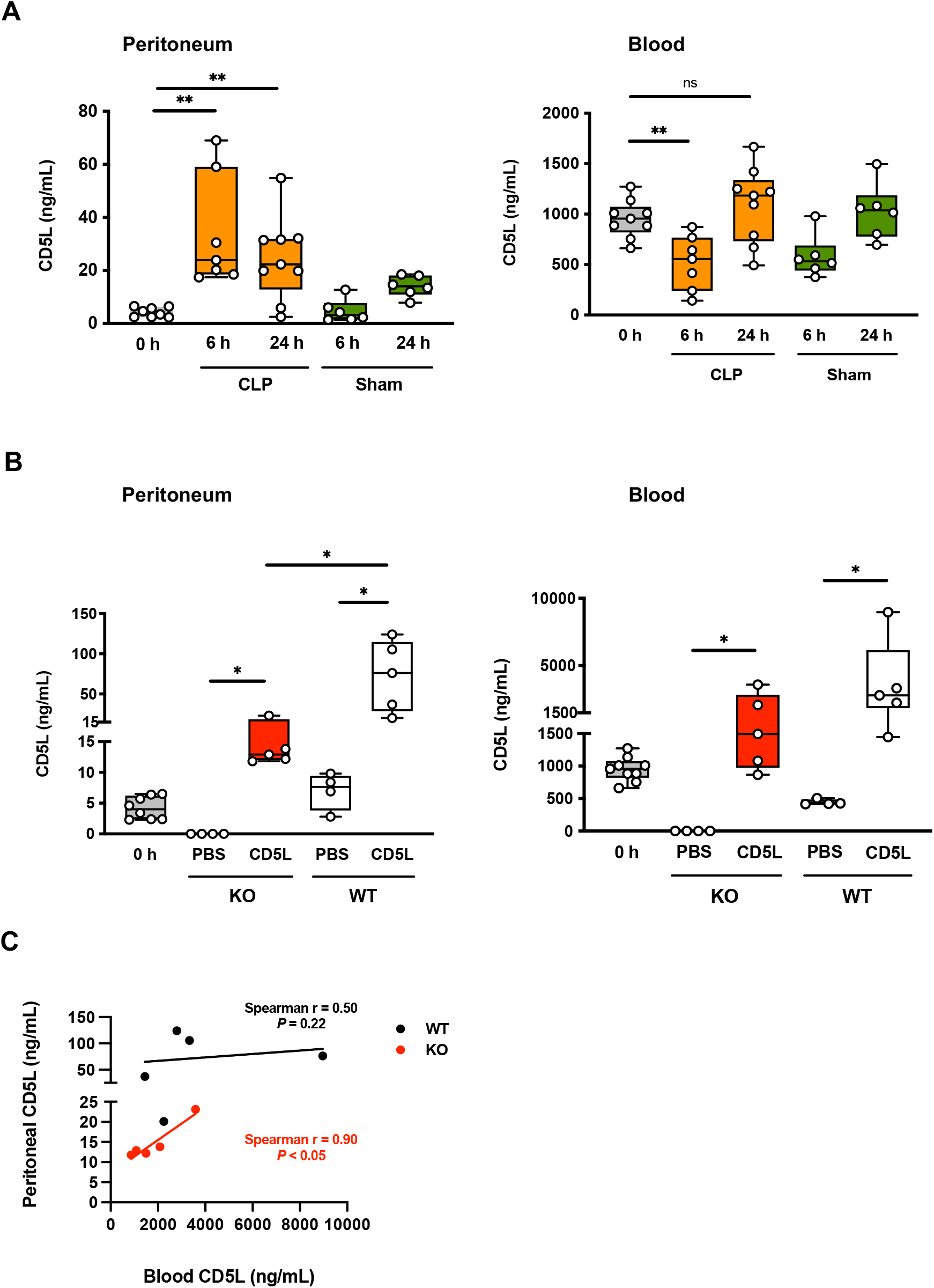
CD5L is trafficked to the site of infection. **(A)** Quantification, by ELISA, of CD5L in the peritoneal cavity (left panel) and serum (right panel) of C57BL/6 mice at 0, 6 or 24 h after mid-grade sepsis (CLP) or in sham operated (sham) animals. **(B)** WT and CD5L-KO naïve healthy mice were subjected to mid-grade CLP and 3 h later were IV administered with recombinant CD5L (rCD5L) at 0 (PBS) or 2.5 mg/Kg (CD5L). Mice were euthanized 1 h after injection to recover the fluids. Amount of CD5L in the peritoneal cavity (left panel) and serum (right panel), quantified by ELISA. **(C)** Scatter graphs with detailed illustration of the relation between peritoneal and blood CD5L with the indication of Spearman’s rank correlation in both WT and CD5L-KO mice upon CD5L IV injection as detailed above. Data from at least 2 independent experiments (Mann-Whitney test). *, *p* < 0.05; **, *p* < 0.01; ns, not statistically significant.

To test the hypothesis of a direct transference of CD5L from the blood to the peritoneum, recombinant mouse CD5L (rCD5L) was injected intravenously (IV) at 2.5 mg/Kg into CD5L-KO mice 3 h after CLP, and peritoneal fluid and blood were recovered 1 h later to quantify the total amount of CD5L by ELISA. One hour after IV administration (4 h after CLP), the amount of CD5L in the blood of the CD5L-KO mice was quantified at ~1-2 μg/ml (**Fig. 3B, right panel**). Importantly, as it confirmed our premise of the direct trafficking between the blood and the infection site, CD5L was clearly detected in the peritoneal fluid (**Fig. 3B, left panel**).

The same procedure was performed on WT mice. and resulted in an even bigger increase of CD5L in the peritoneal cavity, that presumably results from the contribution of both exogenous rCD5L as well as endogenous CD5L. Regarding the levels in the blood, whereas in animals not injected with rCD5L the levels of the endogenous protein dropped, in those mice where rCD5L was IV administered the level of total CD5L was much higher than that seen in CD5L-KO mice, probably again resulting from the contribution of endogenous plus exogenous CD5L. This is reflected in the lack of correlation between peritoneal and blood CD5L in WT mice, contrasting with the statistically significant correlation between the two variables in CD5L-KO mice (**Fig. 3C**).

### Neutrophils from CD5L-KO mice do not have impaired phagocytic or killing capacity

CD5L has been described as a PRR capable to bind to bacteria and, in particular *in vitro* models, to increase the phagocytosis of particles by macrophages and neutrophils [14–16, 26]. Given the susceptibility of CD5L-KO mice to CLP and the impaired number of neutrophils observed in these mice, we investigated the contribution of CD5L to the phagocytosis mediated by neutrophils. Thioglycolate-elicited neutrophils were recovered from the peritoneal cavity of both WT and CD5L-KO mice 6 h after injection. The cell population obtained was routinely highly enriched in neutrophils, identified by the surface expression of Ly6G and CD11b, without significant differences between WT and KO mice (**Fig. 4A**). Before being added to the neutrophils, pHrodo™ Red *E. coli* BioParticles™ were incubated for 1 h with 10 μg of rCD5L, or PBS (0 μg of rCD5L), and were shown to be efficiently bound by rCD5L (**Fig. 4B**). These particles become fluorescent only in acidic pH solution, allowing to distinguish between the bacteria that are inside the phagosomes from those that stay outside the phagocytes [27]. By gating the chosen leukocyte sub-populations, we could assertively measure phagocytosis by neutrophils only. Peritoneal neutrophils recovered from WT and CD5L-KO mice were put into contact with inactivated pHrodo *E. coli* incubated, or not, with rCD5L. Analyzing the percentage of pHrodo-labeled WT and CD5L-KO neutrophils, there were no differences between all groups of cells **(Fig. 4C)**. Moreover, the mean fluorescence intensity (MFI), indicative of the number of phagocyted particles, was not different between WT or CD5L-KO neutrophils, having these been exposed either to rCD5L-pre-coated or to naked pHrodo *E. coli* particles **(Fig. 4D)**.

**Figure 4.**
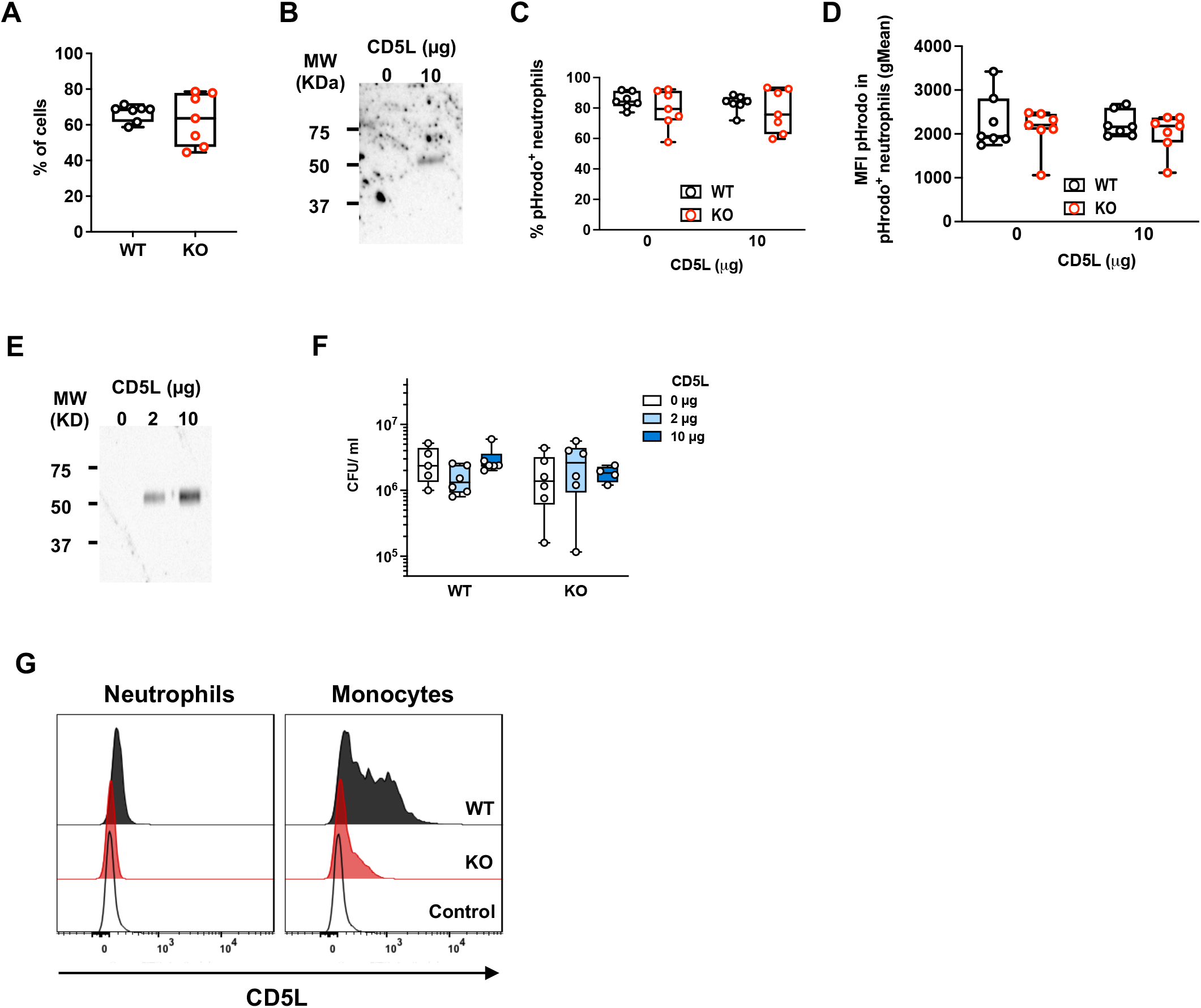
Neutrophils from CD5L-KO mouse do not display defective phagocytosis. **(A)** Percentage of thioglycolate-elicited Ly6G^+^CD11b^+^ neutrophils among all cells collected from the peritoneal cavity of WT and CD5L-KO mice, determined by flow cytometry. **(B)** pHrodo™ Red *E. coli* BioParticles™ were incubated with 0 or 10 μg of rCD5L for 1 h on ice in TBS containing Ca^2+^. After washing, samples were resuspended in Laemmli buffer and heated at 95 °C for 10 min. Western blot detection of CD5L bound to the particles, using mouse anti-His mAb followed by a secondary goat anti-mouse HRP antibody. **(C-D)** Phagocytosis of pHrodo particles. Neutrophils were cultured in the presence of pHrodo particles pre-coated with 0 or 10 μg of rCD5L. **(C)** Percentage of Ly6G^+^CD11b^+^ neutrophils that internalized pHrodo particles, quantified by flow cytometry. **(D)** MFI values (geometric mean) of pHrodo channel within the pHrodo^+^ neutrophil population. **(E)** Cecal bacteria were incubated with 0, 2 and 10 μg of rCD5L for 1 h on ice in TBS containing Ca^2+^. Detection of bacteria-bound rCD5L as in (B). **(F)** Neutrophils from WT or CD5L-KO mice were cultured in the presence of cecal bacteria pre-coated with 0, 2 or 10 μg of rCD5L. After 3 h, cells were washed, lysed and plated for CFU enumeration. **(G)** Intracellular staining of CD5L in WT or CD5L-KO naïve mouse blood neutrophils (Ly6G^+^CD11b^+^) and monocytes (F4/80^+^CD11b^+^) was performed after fixation and permeabilization, with goat anti-mCD5L polyclonal antibody followed by Alexa 488-coupled donkey anti-goat secondary antibody and analyzed by flow cytometry. Data from at least 2 independent experiments. Mann-Whitney test did not reveal any statistical difference between relevant groups.

We additionally performed killing assays to assess the capacity of WT or CD5L-KO neutrophils to destroy phagocyted bacteria, either naked or coated with rCD5L. Thioglycolate-elicited neutrophils recovered from the peritoneal cavity of WT or CD5L-KO mice were cultured with bacteria extracted from mouse cecum, previously incubated and bound with 2 or 10 μg of rCD5L, or naked (0 μg of rCD5L) (**Fig. 4E**). After 3 h of culture, neutrophils were recovered and lysed, and CFU counts indicated that WT or CD5L-KO neutrophils had equivalent killing efficiency and that precoating the bacteria with exogenous rCD5L, at different amounts, had no influence on the killing ability of either population of neutrophils (**Fig. 4F**). Of note, contrasting to monocytes, neutrophils to not produce CD5L (**Fig. 4G**).

### CD5L administration prevents lethality upon CLP

Given that the absence of CD5L increased the lethality in the CLP model, and that there was a significant increase of this protein in WT animals upon infection induced by the surgery, we investigated whether the exogenous administration of the recombinant protein could influence the outcome of sepsis. WT C57BL/6 mice were subjected to CLP to induce high-grade sepsis, following which a group received sequential doses, injected intraperitoneally, of 2.5 mg/Kg of rCD5L after 3 and 6 h, while the control group received PBS alone at the same time points. Survival was monitored for 10 days, and whereas untreated mice had a low survival rate (17%), the percentage of survival of the animals treated with rCD5L was 55% **(Fig. 5A)**. Distributing the dosage of rCD5L through four doses of 1.25 mg/Kg administered at 1, 3, 6 and 24 h resulted in a similar survival rate (62%).

**Figure 5.**
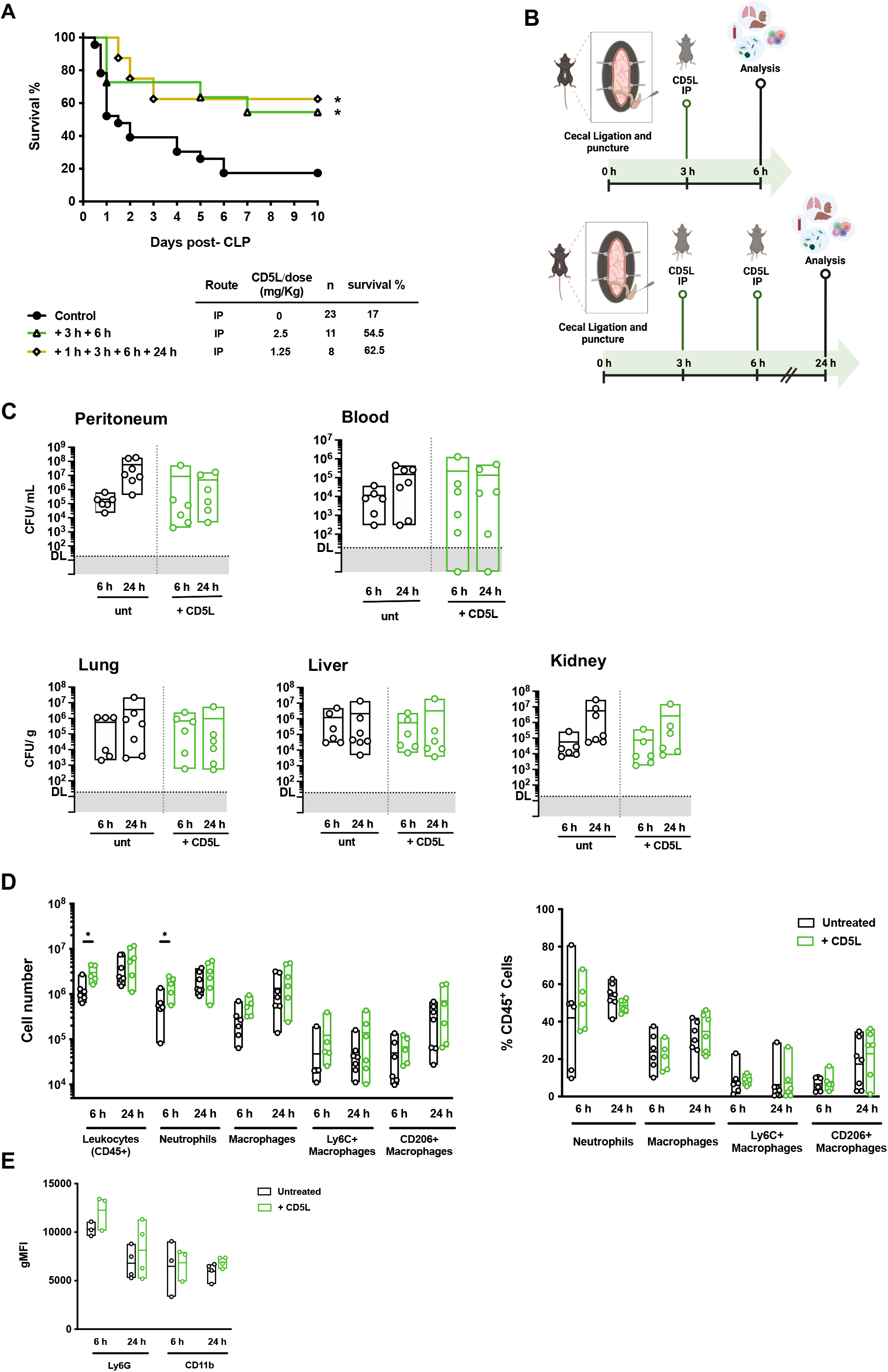
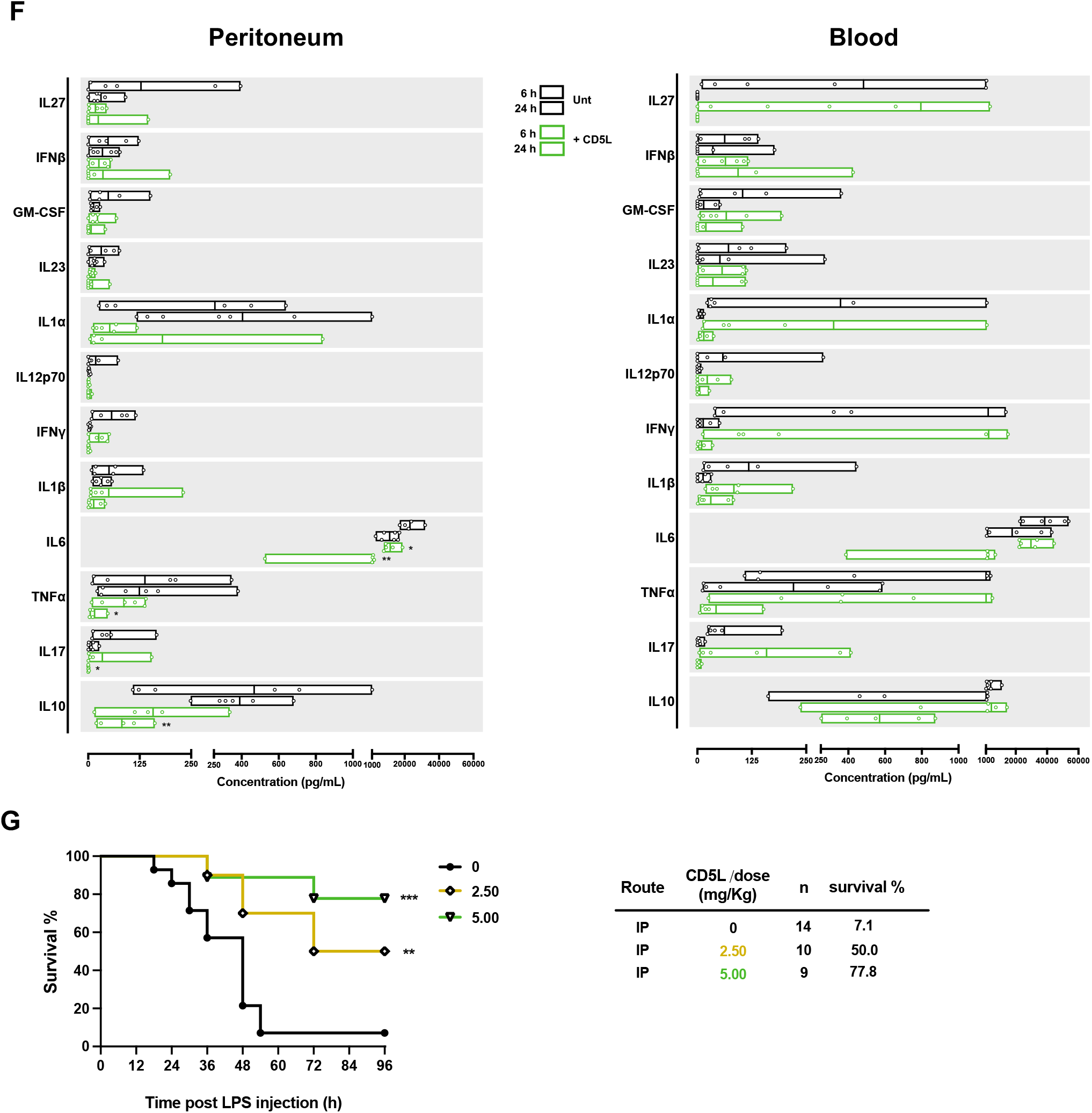
CD5L administration reduces lethality upon sepsis induction. **(A)** WT C57BL/6 mice were subjected to CLP to induce high-grade sepsis. For that, animals were subjected to surgery, exposure and ligation of the cecum (corresponding to approximately 75% the distance between the distal pole and the base of the cecum) and through-and-through puncture with a 21G needle. Upon CLP, mice were injected IP at the indicated times (after surgery) and doses of rCD5L. Kaplan–Meier survival curves were generated to compare mortality between the treated and untreated groups and significance was determined by log-rank (Mantel-Cox) test. The number of mice per group is indicated (n). **(B)** Protocol for analysis of the immune response after CLP and rCD5L IP treatment. Top, analysis at 6 h, mice had been injected IP with 2.5 mg/Kg rCD5L at 3 h post-surgery, or PBS (untreated), and euthanized 3 h later. Bottom, analysis at 24 h, mice were IP injected with 2 doses of 2.5 mg/Kg rCD5L at 3 and 6 h after surgery, or PBS (untreated), followed by euthanasia at 24 h. **(C)** CFU counts of bacteria obtained from the peritoneal cavity, blood, lung, liver and kidney. **(D)** Absolute counts and frequencies of cellular populations in the peritoneal cavity, measured by flow cytometry. Subsets analyzed: leukocytes, CD45^+^; neutrophils, CD45^+^CD11b^+^Ly6G^+^; macrophages, CD45^+^CD11b^+^CD11c^-^F4/80^+^. Ly6C and CD206 markers were used to distinguish between M1 and M2 within the macrophage population. **(E)** Ly6G and CD11b MFI in expressing neutrophils, obtained by geometric mean statistics. **(F)** The indicated cytokines were quantified by a multiplex bead-based immunoassay in samples from the peritoneal cavity (left panel) and blood serum (right panel). **(G)** WT C57BL/6 mice were IP injected with a lethal dose (10 mg/Kg) of LPS and 3 h later with 2.50 or 5 mg/Kg rCD5L, or left untreated (0). Kaplan-Meier survival analysis comparing treated with untreated groups. The number of mice per group is indicated (n). Data from at least 2 independent experiments (Mann-Whitney test). *, *p* < 0.05; **, *p*< 0.01; ***, *p* < 0.005.

To understand the molecular mechanisms involved in CD5L-mediated protection, the progression of infection and the immune response were analyzed: i) at 6 h after CLP in mice that had been treated with a single dose of 2.5 mg/Kg rCD5L given intraperitoneally 3 h post-CLP; and ii) at 24 h in mice that had received two doses of 2.5 mg/Kg, at 3 and 6 h after CLP (**Fig. 5B)**. Bacterial counts were measured locally (peritoneal cavity) and systemically (blood, lungs, liver, kidneys), and although the CFUs showed a tendency to be lower at 24 h in the treated mice, the differences were not significant (**Fig. 5C**).

A detailed analysis of the infiltrating cells in the peritoneal cavity showed that rCD5L IP treatment resulted in an increase in the number of neutrophils recruited at the earlier time point, although without changes in cellular frequencies (**Fig. 5D**). However, the activation phenotype of the recruited neutrophils was not different comparing treated with untreated mice (**Fig. 5E**). IP injection of rCD5L resulted also in substantially reduced amounts of the inflammatory mediator IL-6 at both 6 h and 24 h post CLP in the peritoneal cavity of treated animals, compared with the untreated group. Other inflammatory mediators like TNF-α and IL-17 and the antiinflammatory cytokine IL-10 followed the same pattern at 24 h after infection in the peritoneum of treated mice (**Fig. 5F**). However, no differences in cytokines were detected, between treated and untreated groups, in the blood of both groups of mice during treatment (**Fig. 5F).**

CD5L has been described as also having an anti-inflammatory potential, besides the PRR properties [12, 16, 28]. Given that in the CLP model the anti-inflammatory component seemed to play the major role, we investigated the potential curative role of rCD5L in the model of sterile sepsis. WT C57BL/6 mice received a lethal dose (10 mg/Kg) of LPS, and 2 groups of mice received doses of 2.5 or 5 mg/Kg of rCD5L, given IP 3 h after the LPS challenge, while a third group received PBS only (untreated). Strikingly, whereas the majority of untreated mice died within 2.5 days, mice having received rCD5L at different doses had a survival rate of 50% or more, four days after the LPS challenge (**Fig. 5G).**

### Intravenous therapeutic administration of rCD5L promotes bacterial clearance, reduces inflammation, and significantly increases survival upon polymicrobial infection and sepsis

From the analysis of immune responses of WT mice to mid-grade CLP, and of WT mice treated with rCD5L administered IP after high-grade CLP induction, both models showing a high level of survival, a discordant parameter was that peritoneal recruited neutrophils had acquired a more marked activated phenotype in WT comparing with CD5L-KO mice in the former situation, but not in the latter comparing treated and untreated groups. We thus hypothesized that CD5L would not only help on the mobilization of neutrophils, but that circulatory CD5L could also have a role in driving neutrophils to the activated state. Therefore, we evaluated the possible therapeutic efficacy of rCD5L in fighting CLP-induced sepsis once administered via the IV route.

We first determined the half-life of rCD5L in circulation, by IV administration of the protein in CD5L-KO mice, to be approximately 9 h (**Fig. 6A**). We next went on to analyze the effect of rCD5L IV administration on the recruitment of immune cells to the peritoneum following high-grade CLP induction in WT C57BL/6 mice, and according to the same experimental scheme as shown for the IP treatment **(Fig. 6B)**. IV treatment with rCD5L was characterized by a fast recruitment of neutrophils, both in number as well in percentage, to the site of infection **(Fig. 6C)**. Importantly, the recruited neutrophils displayed a more pronounced activated phenotype comparing with untreated mice, in agreement with our hypothesis **(Fig. 6D)**. As result of the fast and efficient response following rCD5L administration, there was a significant reduction in the bacterial loads in the lungs and liver, and also a small but not significant decrease in the kidney, peritoneum and blood, comparing rCD5L-treated mice with untreated animals **(Fig. 6E)**.

**Figure 6.**
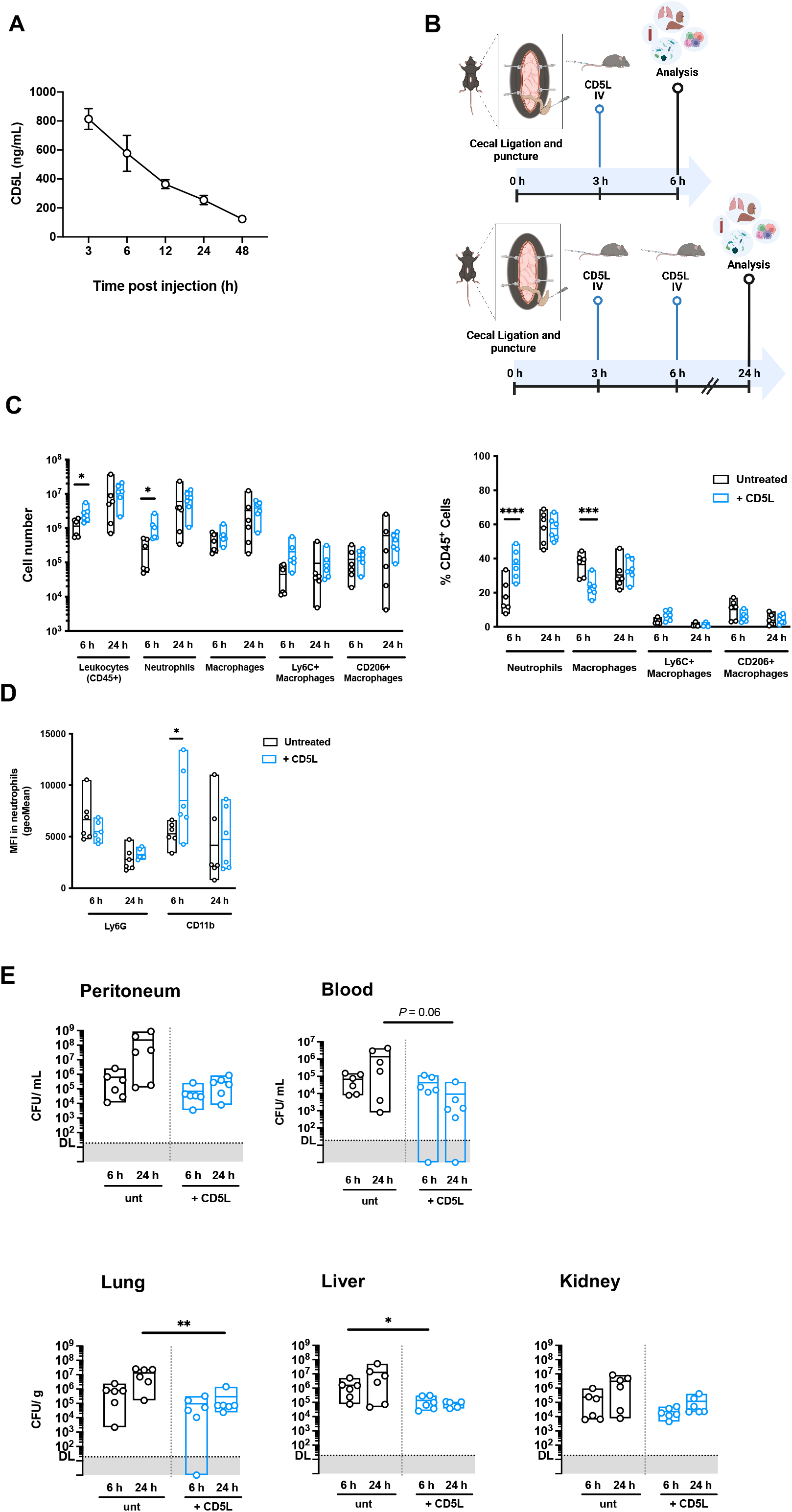
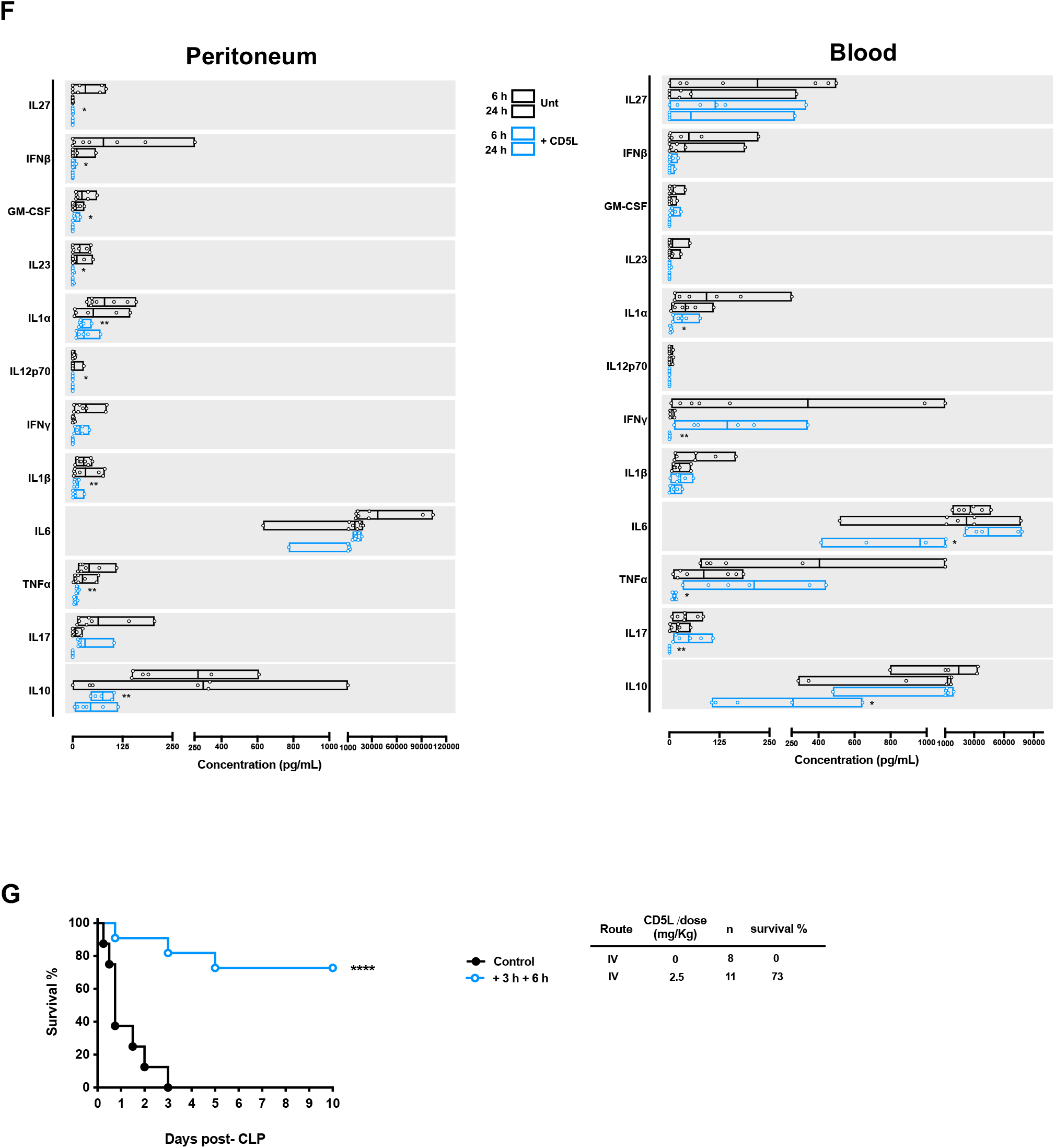
rCD5L IV administration promotes bacterial clearance, reduces inflammation, and prevents lethality upon CLP and sepsis. **(A)** CD5L-KO mice were IV injected with 2.5 mg/Kg of rCD5L, and the circulating amount of rCD5L was quantified by ELISA at the indicated time-points. **(B)** Protocol for analysis of the immune response after CLP and rCD5L IV treatment. Top, analysis at 6 h, mice had been injected IV with 2.5 mg/Kg rCD5L at 3 h post-surgery, or PBS (untreated), and euthanized 3 h later. Bottom, analysis at 24 h, mice were IV injected with 2 doses of 2.5 mg/Kg rCD5L at 3 and 6 h after surgery, or PBS (untreated), followed by euthanasia at 24 h. **(C)** Absolute counts and frequencies of cellular populations in the peritoneal cavity, measured by flow cytometry. Subsets analyzed: leukocytes, CD45^+^; neutrophils, CD45^+^CD11b^+^Ly6G^+^; macrophages, CD45^+^CD11b^+^CD11c^-^F4/80^+^. Ly6C and CD206 markers were used to distinguish between M1 and M2 within the macrophage population. **(D)** Ly6G and CD11b MFI in expressing neutrophils, obtained by geometric mean statistics. **(E)** CFU counts of bacteria obtained from the peritoneal cavity and blood, grown in aerobic or anaerobic conditions, or from lung, liver and kidney, grown in aerobiosis. **(F)** The indicated cytokines were quantified by a multiplex bead-based immunoassay in samples from the peritoneal cavity (upper panels) and blood serum (lower panels). Data from at least 2 independent experiments (Mann-Whitney test). **(G)** C57BL/6 mice were subjected to CLP to induce highgrade sepsis. At 3 and 6 h after the surgery, mice were injected IV with doses of 2.5 mg/Kg of rCD5L. Kaplan–Meier survival curves were generated to compare mortality between the two groups and significance was determined by log-rank (Mantel-Cox) test. The number of mice per group is indicated (n). *, *p* < 0.05; ****, *p* < 0.0001.

The reduction of some inflammatory mediators in the peritoneal cavity after rCD5L IV administration was much faster than that seen for IP-treated animals **(Fig. 5F)**, as one single dose of rCD5L reduced the levels of the inflammatory cytokines IL-27, IFN-β, GM-CSF, IL-23, IL-1α, IL-12p70, IL-1β and TNF-α, already 3 h after treatment (6 h after CLP) **(Fig. 6F)**. The anti-inflammatory mediator IL-10 was also decreased faster upon IV than IP treatment. Systemically, the CD5L IV-treated group had significant changes in the levels of the inflammatory cytokines IL-6, TNF-α and IL-17 and the anti-inflammatory IL-10 at 24 h post CLP when compared with untreated animals (**Fig. 6F, right panel**).

Having characterized the biological and immunological parameters following the IV administration of rCD5L, we followed suit to test the efficacy of rCD5L IV therapy to fight CLP-induced sepsis. WT C57BL/6 mice were submitted to the lethal CLP procedure, and a group of mice received two doses of 2.5 mg/Kg of rCD5L injected IV at 3 and 6 h after CLP, while another group received vehicle alone at the same time points. As seen in **Fig. 6G**, this therapeutic procedure was very effective as the treated animals showed a survival rate of 73%, compared with no survival whatsoever (0%) for the mice receiving no treatment.

### CD5L is associated with increased CXCL1 levels responsible for accentuated neutrophil chemotaxis

Given that the deficiency in CD5L in the mutant mice resulted in decreased immune cell recruitment, and that by opposition the therapeutic administration of rCD5L increased the leukocyte infiltration into the peritoneum, we hypothesized that the protein could be shaping the chemokine profile elicited upon CLP. We thus performed an inflammatory chemokine array in the fluids from the peritoneal cavity and blood of WT and CD5L-KO mice, recovered 3 and 6 h after CLP (**Fig. S3A**), and from mice treated IV with rCD5L 3 h after CLP and euthanized at 6 h (**Fig. S3B**). In both experimental setups, there were undefined variations depending on the analyzed site and mouse group. However, CXCL1, a chemokine connoted with neutrophil chemotaxis, was consistently decreased in the peritoneal cavity and blood of CD5L-KO mice, when compared with WT mice, at 6 h after CLP (**Fig. 7A**). Conversely, mice that were IV treated with rCD5L had significantly higher blood levels of CXCL1 at 6 h after CLP than those untreated (**Fig. 7B**), at the exact same time point where the major difference in neutrophil recruitment in these groups was observed (**Fig. 6C**). These results highlight a novel role for CD5L in controlling chemokine profiles elicited upon an infection, particularly of CXCL1 in this case, that can influence the extent of leukocyte recruitment and lead to enhanced and faster immune responses and ultimately to increased survival.

**Figure 7.**
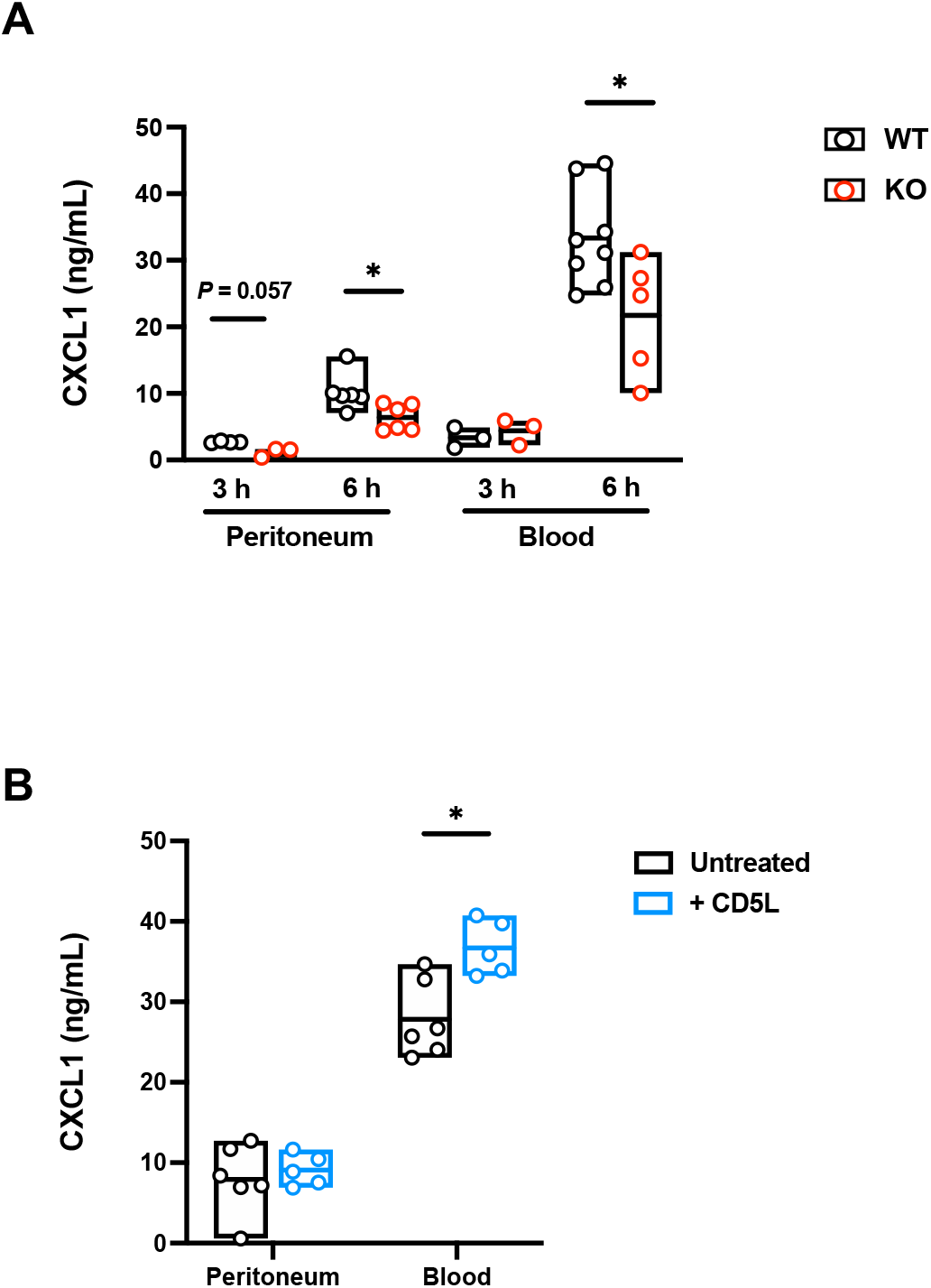
CD5L is associated with the regulation of neutrophil-chemotactic CXCL1 levels. **(A)** WT and CD5L-KO mice were subjected to CLP to induce mid-grade sepsis, and CXCL1 was quantified in the peritoneal cavity or blood serum 3 h or 6 h after surgery. **(B)** WT C57BL/6 mice were subjected to CLP to induce high grade sepsis and IV injected with 2.5 mg/Kg of rCD5L, or with PBS (untreated) 3 h after surgery. CXCL1 was quantified by a multiplex bead-based immunoassay in the peritoneal cavity and blood serum 6 h after CLP. Data from at least 2 independent experiments (Mann-Whitney test). *, *p* < 0.05.

## Discussion

With an estimated 49 million cases and 11 million deaths reported in the year 2017, sepsis is the direct cause of nearly 20% of all deaths worldwide [29]. However, currently there is no treatment against sepsis that shows high efficacy, and the improvements in the last decades have been minimal. Therapies with endotoxin antagonists, immunoglobulins, high- or low-dose corticosteroids, anti-inflammatory cytokine antagonists, anti-TNF antibodies, anticoagulants, etc., all have been largely unsuccessful against sepsis and/or septicemia, many times with more adverse effects than the little benefits achieved. In this study, we basically explored the increase in the bioavailability of the naturally occurring circulating protein CD5L to fight a very deleterious inflammatory syndrome. It carries minimal toxicological effects and has given proofs of anti-inflammatory efficacy, implicating a reduced time frame for approval as an agent for human therapy.

CD5L is present in the blood at relevant concentrations, ~5 μg/ml [30, 31], but it is reported that in several inflammatory contexts, like type 2 diabetes, liver fibrosis, asthma, atopic dermatitis, infection, and sepsis, there is a significant increase in the circulating levels of CD5L [28, 32]. It is disputed whether this augment is in itself an aggravating pathological factor, or whether it represents the increase of a protective factor that is overproduced to fight the causes of disease. The fact that our CD5L-KO mice are more susceptible to a mild controllable form of the inflammatory syndrome seems to support the latter hypothesis. Although not yet fully explored, the functions of CD5L that we disclose involve the recruitment and activation of neutrophils, possibly through the induction of the CXCL1 chemokine by a still undefined mechanism. Adding to this novel property of CD5L its already established functions as PRR and anti-inflammatory mediator, CD5L could thus be considered a multifunctional factor that is able to identify and neutralize an invading microbe, promote neutrophil recruitment, and at the same time contain the infection-associated inflammation.

In accordance with these premises, we showed that mice subjected to lethal-grade CLP treated with rCD5L had an extraordinary rate of survival to this very deleterious procedure, as confirmed by the comparison with the untreated group where virtually all mice succumbed to the disease. This could unequivocally grant to rCD5L a huge therapeutic potential to treat sepsis. However, Gao *et al*. have recently reported that the administration of recombinant mouse CD5L in the CLP model resulted in augmented inflammation, increased bacteremia, and worsened the mortality of the animals [24], casting doubts on the putative anti-inflammatory properties of the protein [24]. It is obviously possible to cross-examine the different methods and procedures used by diverse research teams in order to identify the causes for the completely opposed outcomes of the intended therapy, namely the source of the recombinant proteins, the route of administration, or the different dosages used. Gao and colleagues in fact used doses of rCD5L between 25 and 100-fold lower than those we administered. Nevertheless, it is possible that the different conclusions presented by both groups represent reliable results.

A major difference between our procedure and that of the group from Chongqing is that they give the recombinant protein at the time of inducing sepsis by CLP, whereas we include a delay in the treatment for 3 or more hours after CLP. We do so because, with a view to translate the treatment to the human setting, it may not be useful or instructive to administer a drug at the exact time that the disease is provoked. The therapeutic protocol that we use takes into consideration that the treatment for sepsis should be applied when the disease develops, and thus we contemplate a delay in the first administration of therapeutic rCD5L. A simple explanation to the apparent conflicting results is that, as stated by the authors, the key molecular mechanism to the apparent detrimental action of CD5L is the induction of the anti-inflammatory cytokine IL-10, an observation made previously in other reports [16, 28]. If we consider that CD5L is an immunosuppressant, as deduced by the results obtained from our CD5L-KO mouse model, Gao *et al*., by administering CD5L at the same time of disease initiation, are suppressing the natural response to the initial inflammation, possibly through the induction of elevated IL-10 levels that are known to be detrimental in the early phases of sepsis [33], contrasting with a beneficial role in later phases.

These discussions serve, nonetheless, to highlight the extreme caution that should be given in considering future therapies to treat sepsis, namely the timing of therapeutic intervention. It is not unusual that many successful protocols obtained from animal models cannot in fact be used in the translation to the clinic because in the experimental models very often the therapeutic drug is administered prophylactically or at the time of disease induction. Taking these aspects into consideration and trying as much as possible to simulate the clinical setting, thus contemplating acceptable delays in the diagnostic, transport of patients and other variables, and applying the initial therapy once the disease is diagnosed, our protocol considers the start of the therapeutic procedure 3 hours after the induction of the disease. Remarkably, we obtain a rate of survival above 70%, compared with no survival whatsoever for the untreated animals, in the absence of antibiotics or other complementary drugs. We trust that the noticeable benefits obtained in our animal models can quickly be exploited and translated to clinical medicine.

## Methods

### Development of CD5L KO mice and immunophenotyping

All experiments were conducted in 8- to 12-week-old C57BL/6 mice following the Portuguese (Portaria 1005/92) and European (Directive 2010/63/EU) legislations concerning housing, husbandry and welfare. The project was reviewed and approved by the Ethics Committee of the Instituto de Investigação e Inovação em Saúde (i3S), Universidade do Porto, and by the Portuguese National Entity Direcção Geral de Alimentação e Veterinária (license reference: 009951).

CD5L-knockout (KO) mice were generated using the CRISPR/Cas9 system, as described previously [34]. In brief, a sgRNA (10 ng/μl), targeting the sequence 5’-GGCTGATATGATGCGCCACG-3’, located in exon 3 of *Cd5l* gene, Cas9 mRNA (20 ng/μl), and the replacement ssDNA oligonucleotide containing the 60-nucleotide sequences on each side of the deletion separated by an *Eco*RI site and three tandem stop codons 5’-GGGACCGGCGTGATGTGGCTGTGGTGTGCCGAGAGCTCAATTGTGGAGCAGTCATCCAAACCCCGCGTTAATGATAGGATCCGGCGCATCATATCAGCCACCAGCATCAGAGCAAAGAGTTCTTATTCAAGGGGTTGACTGCAACGGAACGGAAGAC-3’ (10 ng/μl) (obtained from IDT), in 10 mM Tris-HCl pH 7.5, 0.1 mM EDTA were introduced by pronuclear injection of C57BL/6 fertilized oocytes using standard procedures [35]. The sgRNAs were generated by annealing oligos 5’-AGGGGGCTGATATGATGCGCCACG-3’ with 5’-AAACCGTGGCGCATCATATCAGCC-3’, and introducing them into the plasmid gRNA basic [34]. The plasmids were linearized and the sgRNAs synthesized by *in vitro* transcription (Megashort Script in vitro T7, Ambion), purified (Megaclear RNA kit, Ambion) and analyzed on Experion (Bio-Rad) for integrity, purity and yield check. Deletions were assessed by PCR from tail genomic DNA and confirmed by direct sequencing.

In-depth immune-phenotyping analysis was performed on spleens, thyme, peripheral blood and peritoneal cavity of six 12-week-old C57BL/6/N and CD5L-KO mice (3 males, 3 females per group) by multi-parametric flow cytometry. Leukocytes were isolated from thymi and spleens by combined mechanical treatment and enzymatic digestion using a GentleMACS™ octo dissociator system (Miltenyi). Cell suspensions were numerated for all samples on AttuneNxT flow cytometer. Erythrocyte were lyzed using RBC Lysis solution (eBioscience, #00-4333-57) following manufacturer recommendations. Fc-receptors were blocked using anti-CD16/CD32 (#24G2) hybridoma. Cells were stained in one step with corresponding antibody panel (**Suppl. Table 1 and 2**), washed and analyzed within 4 h after organ collection. Dead cells were removed using Live/Dead SytoxBlue staining (Thermo, #S11348). Datasets were acquired on a 5-laser BD LSRFortessa™ SORP cell analyzer using an HTS plate reader on a standardized analysis matrix using BD FACSDiva 8.0.1 software. Data were visualized using radar plot on Excel or Tableau Desktop software. Frequencies of cell subsets on all mice of the study were pooled in one single spreadsheet per panel per organ, transformed in asinh and centered to the mean. The resulting variable list was run in SIMCA Multivariate analysis software (Sartorius). First, distribution of samples in PCA score was considered to remove potential outliers. Next, an OPLS-DA method was applied on the remaining dataset to identify groups of samples presenting similar types of variations. Overall, the distribution of mice was similar before and after OPLS-DA modelling for each experimental group. Variables important for this projection (VIP) were selected with a VIPpred value over 1. A final PCA was run on data only coming from these selected predictive variables to ensure that the OPLS-DA model was not overfitting the dataset. List of phenotypes with VIP > 1 were further visualized on moustache plots using Tableau Desktop software.

### CD5L cloning, production and purification

Mouse CD5L cloning, production and purification was performed by INVIGATE. Briefly, mouse *Cd5l* cDNA was amplified from murine cDNA extracts and cloned into vector pcDNA3.1/Zeo(+) together with a C-terminally fused sequence coding for an 8x histidine tag. For efficient secretory expression, the *mCd5l* cDNA was placed downstream of the coding sequence for the signal peptide of human CD14 protein (Met1-Val17). The resulting plasmid pCD14SP-mCD5L-8HIS_Zeo was transfected in HEK293T cells and the cells were selected by supplementation of 150 μg/ml Zeocin. Cells stably expressing mouse CD5L were grown in roller bottle cultures. Collected culture supernatants were subjected to ultrafiltration (5 KDa Sartocon Slice Cassette) to replace culture medium with 50 mM Tris, 200 mM NaCl pH 7.8 (Tris/NaCl). Sample was purified by immobilized metal affinity chromatography (IMAC) using Ni-NTA-Agarose-Resin (Genaxxon bioscience). Resin was washed with Tris/NaCl/50mM imidazol followed by elution with Tris/NaCl/350 mM Imidazol. Fractions containing mCD5L were identified using SDS-PAGE, pooled and diluted with Tris/NaCl (1:20). For further purification sample was loaded on Q-Sepharose FF (GE Healthcare), washed with Tris/NaCl and eluted with linear gradient to 1 M NaCl. After dialysis and further concentration in centrifugal concentrators the protein concentration was determined by photometric measurement (A280 nm) and silver stained SDS-PAGE was performed to confirm homogeneity of > 95%. Endotoxicity of the 0.22 μm filtered mCD5L-solution was assessed to be less than 1 EU/μg protein by limulus amebocyte lysate chromogenic endpoint assay (Associates of Cape Cod Europe GmbH). The recombinant mouse CD5L protein preparation was aliquoted and lyophilized for convenient storage and use.

### Immunofluorescence

Mouse peritoneal cells were adhered to poly-L-lysine-coated glass slides for 30 min at 37 °C followed by 4% PFA fixation. Surface staining with an anti-mouse anti-F4/80 antibody was performed followed by an AlexaFluor 594-coupled anti-rat secondary antibody. Intracellular staining with a goat anti-CD5L polyclonal antibody followed by an AlexaFluor 488-coupled donkey anti-goat secondary antibody after permeabilization with Triton X100. Dapi was used as a nuclear counterstaining. Cells were visualized in a Leica DMI6000 microscope.

### ELISA

mCD5L quantification was performed using the Mouse CD5L ELISA Pair Set (SinoBiological) following the manufacturer’s instructions.

### Flow cytometry and multiplex assays

For analysis of cellular populations, cells were recovered, washed with flow staining media (FSM, PBS containing 2% FBS) and 1 x 10^6^ cells were stained with fluorochrome-conjugated antibodies against surface markers. Where necessary, cells were fixed with 2% PFA and permeabilized with 0.1% saponin in FSM before intracellular staining with anti-mCD5L. Cells were analyzed in a FACSCanto II flow cytometer (BD) using FACS Diva software.

Mouse cytokine (LEGENDplex™ Mouse Inflammation Panel, Biolegend) and chemokine (LEGENDplex™ Mouse Proinflammatory Chemokine Panel, Biolegend) were quantified in mouse sera or peritoneal fluid following the provided instructions. Samples were acquired in a Accuri C6 (BD) flow cytometer. Data were analyzed in FlowJo software.

### Phagocytosis and killing assays

Preparation of CD5L-coated pHrodo particles: one vial containing 2 mg of particles (pHrodo™ Red *E. coli* BioParticles™ Conjugate for Phagocytosis, ThermoFisher) were reconstituted with 500 μL of TBS with 5 mM CaCL_2_ and sonicated for 5 min. Particles were split into 2 tubes each containing 250 μL of the reconstituted particles: 10 μg of CD5L was added to one of the tubes and the other was left without protein, as control. Protein was incubated with the particles for 1 h at 4°C with rotation followed by washing three times with PBS and resuspending each tube in 250 μL RPMI without phenol red supplemented with 10% FBS and 1% sodium pyruvate. The confirmation of binding of CD5L with the particles was performed by western-blotting. A 30 μl sample of coated particles was mixed with 6x Laemmli sample buffer, denatured for 10 min at 95 °C and loaded on a 10% SDS-polyacrylamide gel (SDS-PAGE) and separated for 45 min at 200 V (Bio-Rad). Samples were transferred to a nitrocellulose membrane (TransBlot Turbo, Bio-Rad). The membrane was blocked with a solution of 5% non-fat milk in Tris-Buffered Saline and 1% Tween (TBS-T) for 1 h at RT, then incubated with the primary antibody anti-His (Qiagen) at a concentration of 1:2500 in a solution of 3% non-fat milk in TBS-T overnight at 4 °C. The membrane was washed three times with TBS-T for 10 min and subjected to the HRP-conjugated secondary goat anti-mouse antibody (Biolegend) in a solution of 3% non-fat milk in TBS-T for 1 h. ECL solution (GE Healthcare) was used to develop a chemiluminescent signal in an X-ray film (GE Healthcare).

Peritoneal neutrophil recovery: six hours after injecting 2.5 ml of 3% thioglycolate IP, mice were euthanized and 5 ml of RPMI medium (HyClone GE Healthcare Life Sciences) with 5% FBS was injected in the mice peritoneum and left for 1 min before recovery by aspiration with a syringe. Neutrophil enrichment was confirmed by flow cytometry as detailed above.

*In vitro* phagocytosis assays: 1 x 10^6^ peritoneal cells were incubated with 2 μl of *E. coli* pHrodo particles for 1 h to allow for phagocytosis to occur followed by two washes with FACS buffer. Cells were then analyzed by flow cytometry as detailed above.

Cecal bacteria isolation, growth and coating with CD5L: bacteria were isolated from WT mice cecum as described previously [36]. Briefly, the cecal content of 3 mice was removed, diluted in PBS and filtered through sterile gauze. The cecal suspension was diluted in brain heart infusion (BHI) medium (BD Biosciences) and incubated at 37 °C for 18 h. Afterwards, the suspension was centrifuged for 15 min at 4000 rpm and washed twice with PBS. The bacterial suspension was then aliquoted, lyophilized and stored at −80°C.

For the killing assays, a lyophilized aliquot was grown for 20 h in 100 ml of BHI medium, at 37°C with agitation. After that, bacteria were washed with sterile PBS twice to achieve an optical density (OD) at 600 nm of 2.0. For coating with CD5L, 1 x 10^8^ bacteria were resuspended in 500 μl of TBS with 5 mM CaCl2, containing 0, 2 or 10 μg of rmCD5L and incubated for 1 h on ice. After, bacteria were washed twice with PBS. The pellet was resuspended in 200 μl PBS and plated in BHI agar plates by serial dilution for CFU determination. The confirmation of binding of CD5L with bacteria was done by Western-Blot as detailed above.

Killing Assays with cecal bacteria: bacteria were incubated with the peritoneal cells at a multiplicity of infection of 5, in incomplete RPMI medium for 3 h followed by washing twice with PBS. Cells were lysed in 500 μL of 0.2% Triton X-100 (Sigma) in PBS. Bacteria were plated in BHI agar plates and CFU number in the suspension was determined through serial dilutions. CFU were counted overnight.

### Cecal-ligation and puncture model

For induction of sepsis by cecal ligation and puncture (CLP), mice were subjected to surgery under isoflurane anesthesia, exposure and ligation of the cecum (corresponding to approximately half the distance between the distal pole and the base of the cecum for mid-grade sepsis, or 75% of the distance for high-grade sepsis) and through-and-through puncture with a 21G needle. After suturing, mice were subcutaneously (sc) injected with warm 0.9% NaCl. Buprenorphine (0.08 mg/ Kg) was administered sc every 12 h until 48 h post-surgery. Mice were monitored twice a day.

### LPS-induced septic shock

For the induction of septic shock, mice were intraperitoneally (IP) injected with 10 mg/Kg lipopolysaccharides (LPS) from *Escherichia coli* O111:B4 (Sigma-Aldrich). Mice were monitored twice a day.

### Bacterial counts

For quantification of bacteria in the indicated organs, 10-fold serial dilutions of the cellular suspensions were plated in Brain Heart Infusion (BHI) agar plates and grown in aerobic conditions for 18 h at 37 °C.

### Histology

Lung, liver and kidney tissue sections were fixed in neutral buffered formaldehyde prior to paraffin embedding. 4 μm sections were stained with hematoxylin and eosin. Pathology scores of lung, liver and kidney were attributed by an independent pathologist in a double blinded analysis. 0: no signs of inflammation, 1: minimal inflammatory signs, 2: Mild inflammation, 3: Moderate to severe inflammation.

## Acknowledgements

This work was funded by National Funds through FCT – Fundação para a Ciência e a Tecnologia, I.P., in the framework of the projects SRecognite Infect-ERA/0003/2015, to A.M.C., and LISBOA-01-0145-FEDER-030254, to M.M., and by CNRS, INSERM, the SRecognite project (ANR-Infect-ERA-2015), and the Investissement d’Avenir program PHENOMIN (French National Infrastructure for mouse Phenogenomics; ANR-10-INBS-07) to B.M. and H.L. R.F.S. and M.S.C. were recipients of PhD studentships from FCT, references SFRH/BD/110691/2015 and SFRH/BD/116791/2016, respectively.

## Notes

### Competing Interest Statement

Alexandre Carmo and Liliana Oliveira and inventors in the patent application (pending) Recombinant Human CD5L protein, active fragments or peptides derived thereof and pharmaceutical composition comprising the recombinant human CD5L protein, active fragments or peptides derived thereof for the treatment of acute infectious diseases, inflammatory diseases and sepsis. PCT/PT2022/050004.

## References

1. Martinez V.G., et al., The conserved scavenger receptor cysteine-rich superfamily in therapy and diagnosis. Pharmacol Rev, 2011. 63(4): p. 967–1000.

2. Bessa Pereira C., et al., The Scavenger Receptor SSc5D Physically Interacts with Bacteria through the SRCR-Containing N-Terminal Domain. Front Immunol, 2016. 7: p. 416.

3. Cardoso M.S., et al., Physical Interactions With Bacteria and Protozoan Parasites Establish the Scavenger Receptor SSC4D as a Broad-Spectrum Pattern Recognition Receptor. Front Immunol, 2021. 12: p. 760770.

4. Lenz L.L., CD5 sweetens lymphocyte responses. Proc Natl Acad Sci U S A, 2009. 106(5): p. 1303–4.

5. Peiser L., et al., The class A macrophage scavenger receptor is a major pattern recognition receptor for Neisseria meningitidis which is independent of lipopolysaccharide and not required for secretory responses. Infect Immun, 2002. 70(10): p. 5346–54.

6. Oliveira M.I., et al., CD6 attenuates early and late signaling events, setting thresholds for T-cell activation. Eur J Immunol, 2012. 42(1): p. 195–205.

7. Tarakhovsky A., et al., A role for CD5 in TCR-mediated signal transduction and thymocyte selection. Science, 1995. 269(5223): p. 535–7.

8. Van Gorp H., P.L. Delputte, and H.J. Nauwynck, Scavenger receptor CD163, a Jack-of-all-trades and potential target for cell-directed therapy. Mol Immunol, 2010. 47(7-8): p. 1650–60.

9. Ojala J.R., et al., Progressive reactive lymphoid connective tissue disease and development of autoantibodies in scavenger receptor A5-deficient mice. Am J Pathol, 2013. 182(5): p. 1681–95.

10. Miyazaki T., et al., Increased susceptibility of thymocytes to apoptosis in mice lacking AIM, a novel murine macrophage-derived soluble factor belonging to the scavenger receptor cysteine-rich domain superfamily. J Exp Med, 1999. 189(2): p. 413–22.

11. Sanchez-Moral L., et al., Multifaceted Roles of CD5L in Infectious and Sterile Inflammation. Int J Mol Sci, 2021. 22(8).

12. Martinez V.G., et al., The macrophage soluble receptor AIM/Api6/CD5L displays a broad pathogen recognition spectrum and is involved in early response to microbial aggression. Cell Mol Immunol, 2014. 11(4): p. 343–54.

13. Sarrias M.R., et al., A role for human Sp alpha as a pattern recognition receptor. J Biol Chem, 2005. 280(42): p. 35391–8.

14. Arai S., et al., Apoptosis inhibitor of macrophage protein enhances intraluminal debris clearance and ameliorates acute kidney injury in mice. Nat Med, 2016. 22(2): p. 183–93.

15. Haruta I., et al., Association of AIM, a novel apoptosis inhibitory factor, with hepatitis via supporting macrophage survival and enhancing phagocytotic function of macrophages. J Biol Chem, 2001. 276(25): p. 22910–4.

16. Sanjurjo L., et al., CD5L Promotes M2 Macrophage Polarization through Autophagy-Mediated Upregulation of ID3. Front Immunol, 2018. 9: p. 480.

17. Maehara N., et al., AIM/CD5L attenuates DAMPs in the injured brain and thereby ameliorates ischemic stroke. Cell Rep, 2021. 36(11): p. 109693.

18. Kurokawa J., et al., Macrophage-derived AIM is endocytosed into adipocytes and decreases lipid droplets via inhibition of fatty acid synthase activity. Cell Metab, 2010. 11(6): p. 479–92.

19. Tomita T., et al., Apoptosis inhibitor of macrophage ameliorates fungus-induced peritoneal injury model in mice. Sci Rep, 2017. 7(1): p. 6450.

20. Joseph S.B., et al., LXR-dependent gene expression is important for macrophage survival and the innate immune response. Cell, 2004. 119(2): p. 299–309.

21. Sanjurjo L., et al., The scavenger protein apoptosis inhibitor of macrophages (AIM) potentiates the antimicrobial response against Mycobacterium tuberculosis by enhancing autophagy. PLoS One, 2013. 8(11): p. e79670.

22. Wang C., et al., CD5L/AIM Regulates Lipid Biosynthesis and Restrains Th17 Cell Pathogenicity. Cell, 2015. 163(6): p. 1413–27.

23. Wang P., et al., Administration of GDF3 Into Septic Mice Improves Survival. Front Immunol, 2021. 12: p. 647070.

24. Gao X., et al., Therapeutic Targeting of Apoptosis Inhibitor of Macrophage/CD5L in Sepsis. Am J Respir Cell Mol Biol, 2019. 60(3): p. 323–334.

25. Singer M., et al., The Third International Consensus Definitions for Sepsis and Septic Shock (Sepsis-3). JAMA, 2016. 315(8): p. 801–10.

26. Gao X., et al., CD5L contributes to the pathogenesis of methicillin-resistant Staphylococcus aureus-induced pneumonia. Int Immunopharmacol, 2019. 72: p. 40–47.

27. Simons E.R., Measurement of phagocytosis and of the phagosomal environment in polymorphonuclear phagocytes by flow cytometry. Curr Protoc Cytom, 2010. Chapter 9: p. Unit9 31.

28. Sanjurjo L., et al., The human CD5L/AIM-CD36 axis: A novel autophagy inducer in macrophages that modulates inflammatory responses. Autophagy, 2015. 11(3): p. 487–502.

29. Rudd K.E., et al., Global, regional, and national sepsis incidence and mortality, 1990-2017: analysis for the Global Burden of Disease Study. Lancet, 2020. 395(10219): p. 200–211.

30. Desiere F., et al., The PeptideAtlas project. Nucleic Acids Res, 2006. 34(Database issue): p. D655–8.

31. Farrah T., et al., A high-confidence human plasma proteome reference set with estimated concentrations in PeptideAtlas. Mol Cell Proteomics, 2011. 10(9): p. M110 006353.

32. Gao X., et al., Assessment of Apoptosis Inhibitor of Macrophage/CD5L as a Biomarker to Predict Mortality in the Critically Ill With Sepsis. Chest, 2019. 156(4): p. 696–705.

33. Song G.Y., et al., What is the role of interleukin 10 in polymicrobial sepsis: anti-inflammatory agent or immunosuppressant? Surgery, 1999. 126(2): p. 378–83.

34. Casaca A., A. Nóvoa, and M. Mallo, Hoxb6 can interfere with somitogenesis in the posterior embryo through a mechanism independent of its rib-promoting activity. Development, 2016. 143(3): p. 437–48.

35. Hogan B., et al., Manipulating the Mouse Embryo: A Laboratory Manual. 1994: Cold Spring Harbor Laboratory Press.

36. Moreno S.E., et al., IL-12, but not IL-18, is critical to neutrophil activation and resistance to polymicrobial sepsis induced by cecal ligation and puncture. J Immunol, 2006. 177(5): p. 3218–24.

