## Supplemental Figures for "CD5L constraints acute and systemic inflammation and can be a novel potent therapeutic agent against sepsis"

#### Supplemental Figure 1

A

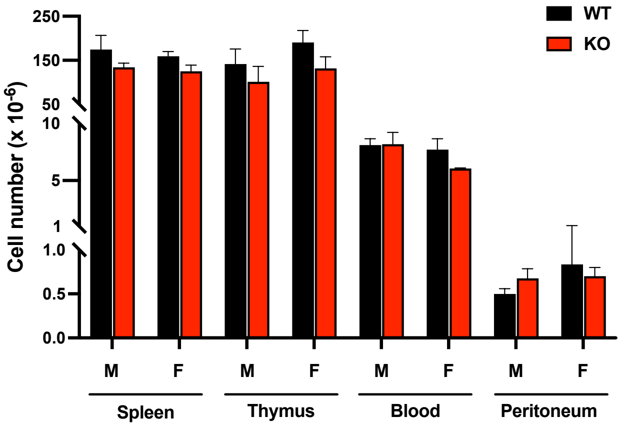


B

*Hematological parameters*

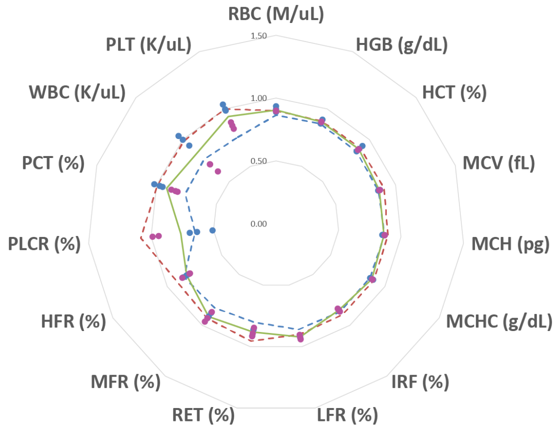


C

*Spleen*

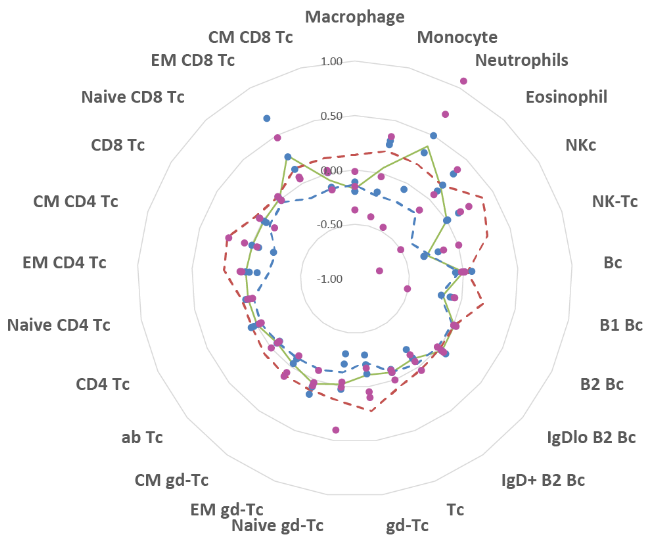

D

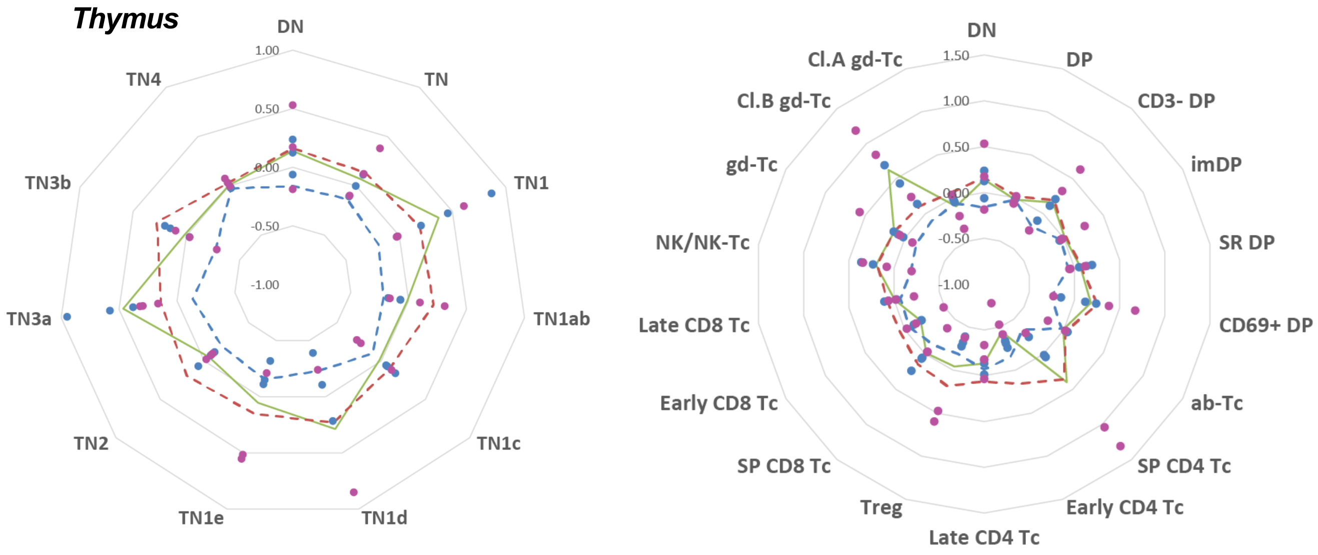


E

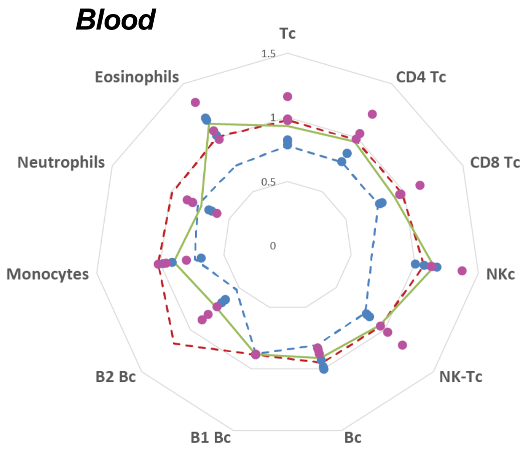


F

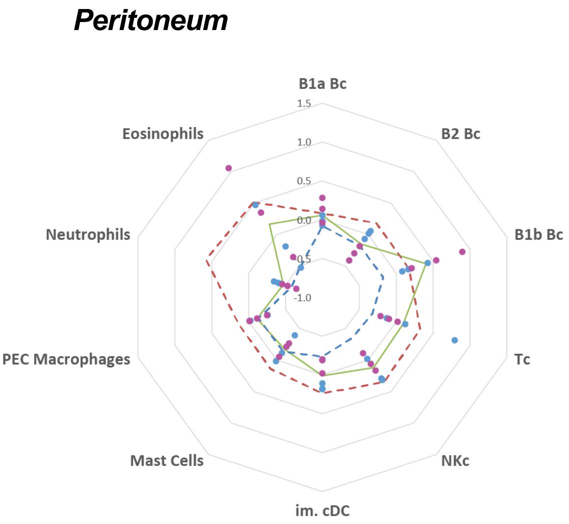


**Figure S1 - Immunophenotyping of CD5L-KO mice. (A)** Total leukocyte numbers in the spleen, thymus, peripheral blood and peritoneal cavity of C57BL/6 WT and CD5L-KO naïve male (M) and female (F) mice. **(B-F)** FC Radar plot of hematological parameters **(B)** and leukocyte proportion variations between CD5L-KO and control animals in spleen **(C)**, thymus **(D)**, blood **(E)** and peritoneal cavity **(F).** EM: effector memory; CM: central memory; TN: triple negative; SP: single positive; DP: double positive; im: immature; ab: αβ; gd: γδ T cells; Bc: B cell, Tc: T cell; NKc: NK cell; PEC: peritoneal resident macrophages. Each circle (blue: male; pink: female) represents a value obtained in independent CD5L-KO mice (value measured in ($KO_n$)/(Mean WT)-1). **Green line**: average variation of subset proportion in CD5L-KO compared with WT animals [(Mean KO)/(Mean WT)-1]; **blue dashed line**: lower limit of WT value dispersion [(Mean WT - 1 SD)/(Mean WT)-1]; **red dashed line:** higher limit of WT value dispersion [(Mean WT + 1 SD)/(Mean WT)-1]. Populations were defined according to the expression of markers detailed in Supplemental table 1. Values expressed in asinh ratio. This high content analysis exemplifies the absence of major leukocyte changes in CD5L-KO mice.

#### Supplemental Figure 2

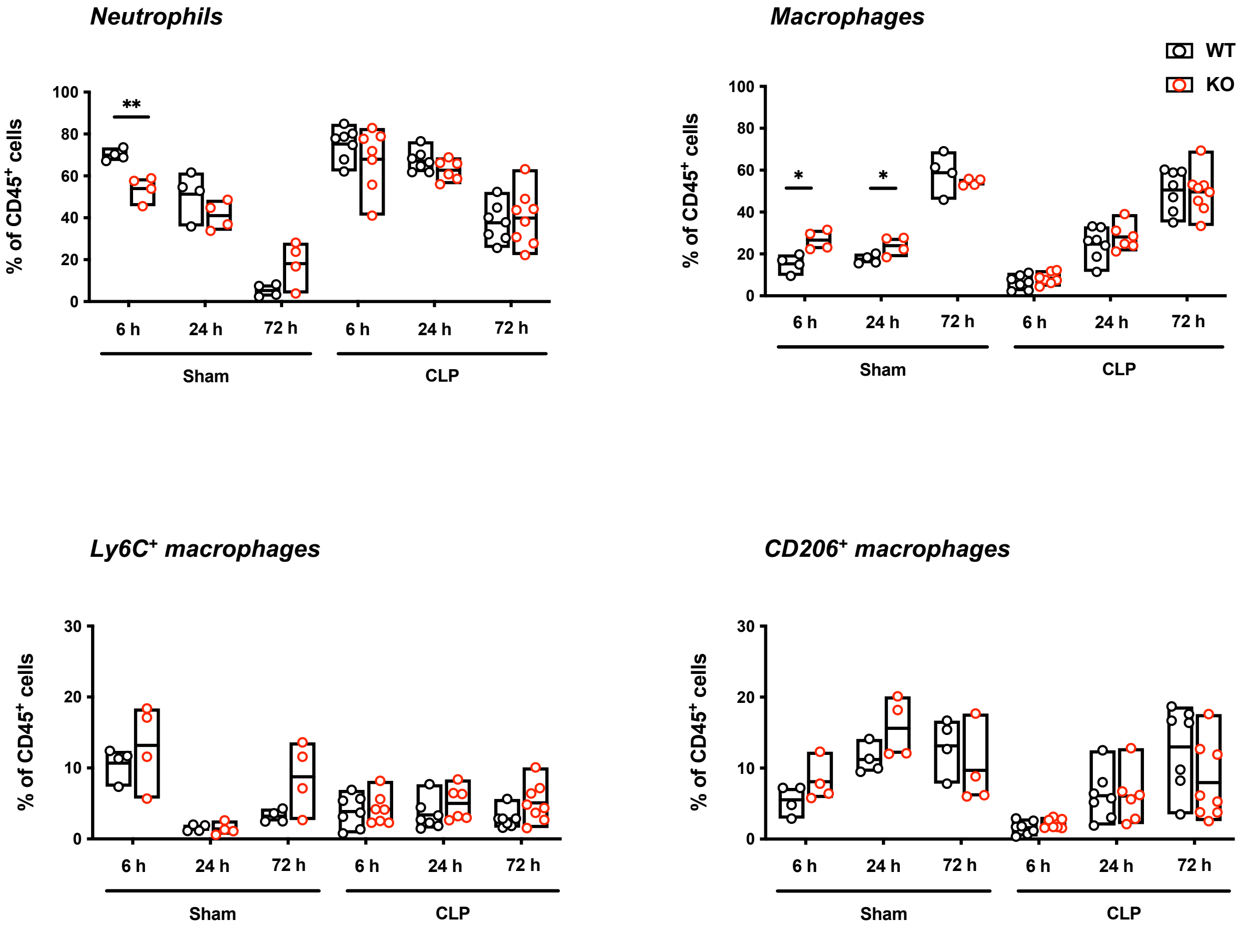


**Figure S2 – Leukocyte sub-populations do not vary between WT and CD5L-KO mice upon CLP, but CD5L-KO mice have an enrichment of neutrophils and a converse decrease in the percentage of macrophages following the placebo surgery.** Frequencies, measured by flow cytometry, of CD45$^+$ cellular populations in the peritoneal cavity of WT and CD5L-KO mice undergoing mid-grade CLP, or sham-operated (subjected to the same surgical procedure but without cecum ligation and puncture). Subsets analyzed: neutrophils, CD45$^+$CD11b$^+$Ly6G$^+$; macrophages, CD45$^+$CD11b$^+$CD11c$^-$F4/80$^+$. Ly6C and CD206 markers were used to distinguish between M1 and M2 within the macrophage population. Data from at least 2 independent experiments (Mann-Whitney test). *, $p < 0.05$; **, $p < 0.01$.

#### Supplemental Figure 3

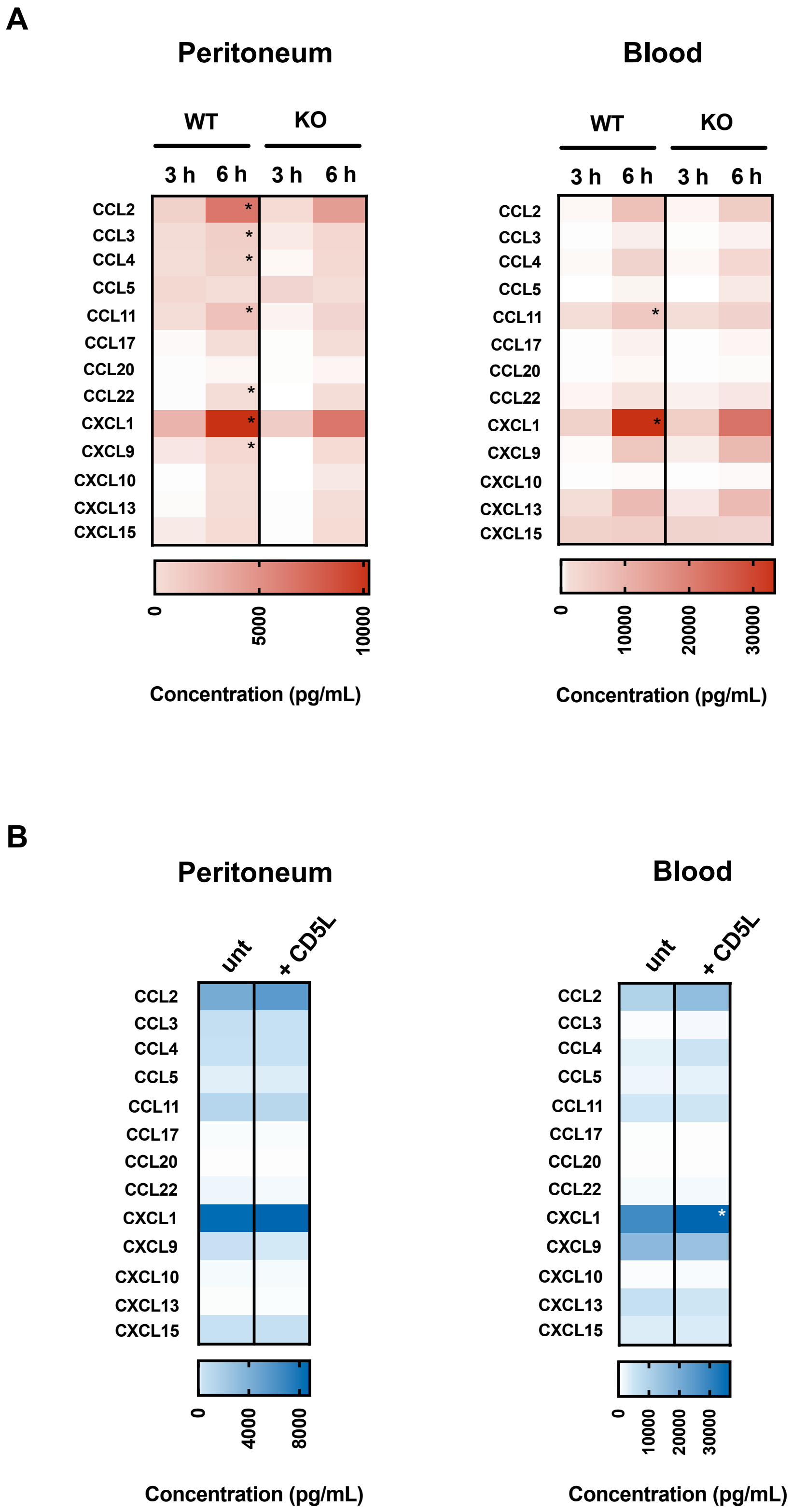


**Figure S3 - CD5L levels correlate with CXCL1 production. (A)** WT or CD5L-KO mice were subjected to CLP to induce mid-grade sepsis, and different inflammatory chemokines were quantified by a multiplex bead-based immunoassay in the peritoneal cavity or blood serum 3 or 6 h after surgery. **(B)** WT C57BL/6 mice were subjected to CLP to induce lethal-grade sepsis, and IV injected with 2.5 mg/Kg of CD5L, or with PBS (untreated), 3 h after surgery. Inflammatory chemokines were quantified in the peritoneal cavity and blood serum 6 h after CLP. Data from at least 2 independent experiments (Mann-Whitney test). *: $p < 0.05$.

**Supplemental Table 1.** Populations described by immunophenotyping panels for spleen, peripheral blood, thymus and peritoneal cavity.

#### Spleen

| Population | Marker Definition | | | | | | | | |
|---|---|---|---|---|---|---|---|---|---|
| **Neutrophils** | CD11b+ | Ly6G+ | | | | | | | |
| **Eosinophils** | CD11b+ | Ly6G- | Ly6Clo | SSChi | | | | | |
| **Macrophages** | CD11bint | Ly6G- | Ly6C- | F4/80hi | CD11c- | | | | |
| **Monocytes** | CD11b+ | Ly6G- | Ly6Chi | | | | | | |
| **pDC** | CD11b- | Ly6G- | Ly6Chi | CD317+ | | | | | |
| **DC** | CD11b- | Ly6G- | Ly6C- | CD317- | CD11c+ | MHCII+ | | | |
| **CD8a-Type DC** | CD11b- | Ly6G- | CD11b+ | CD317- | CD11c+ | MHCII+ | CD8a+ | | |
| **CD11b-Type DC** | CD11b- | Ly6G- | CD11b- | CD317- | CD11c+ | MHCII+ | CD11b+ | | |
| **NKc** | CD161+ | Ly6G- | CD317- | CD5- | | | | | |
| **NK CD11b+Ly6C-** | CD161+ | Ly6G- | CD317- | CD5- | CD11b+ | Ly6C- | | | |
| **NK CD11b+Ly6C+** | CD161+ | Ly6G- | CD317- | CD5- | CD11b+ | Ly6C+ | | | |
| **NK CD11b-Ly6C-** | CD161+ | Ly6G- | CD317- | CD5- | CD11b- | Ly6C- | | | |
| **NK CD11b-Ly6C+** | CD161+ | Ly6G- | CD317- | CD5- | CD11b- | Ly6C+ | | | |
| **NKTc** | CD161+ | Ly6G- | CD317- | CD5+ | | | | | |
| **Bc** | CD161- | Ly6G- | CD317- | CD19+ | MHCII+ | | | | |
| **B1 Bc** | CD161- | Ly6G- | CD317- | CD19+ | MHCII+ | CD5+ | | | |
| **B2 Bc** | CD161- | Ly6G- | CD317- | CD19+ | MHCII+ | CD5- | | | |
| **B2 IgDlo** | CD161- | Ly6G- | CD317- | CD19+ | MHCII+ | CD5- | IgDlo | | |
| **B2 IgD+** | CD161- | Ly6G- | CD317- | CD19+ | MHCII+ | CD5- | IgDhi | | |
| **Tc** | CD161- | Ly6G- | CD317- | CD5+ | CD3+ | | | | |
| **gd-Tc** | CD161- | Ly6G- | CD317- | CD5+ | CD3+ | TCRd+ | | | |
| **EM gd-Tc** | CD161- | Ly6G- | CD317- | CD5+ | CD3+ | TCRd+ | CD44+ | CD62L- | |
| **Naive gd-Tc** | CD161- | Ly6G- | CD317- | CD5+ | CD3+ | TCRd+ | CD44- | CD62L+ | |
| **CM gd-Tc** | CD161- | Ly6G- | CD317- | CD5+ | CD3+ | TCRd+ | CD44+ | CD62L+ | |
| **ab Tc** | CD161- | Ly6G- | CD317- | CD5+ | CD3+ | TCRd- | | | |
| **CD4 Tc** | CD161- | Ly6G- | CD317- | CD5+ | CD3+ | TCRd- | CD4+ | | |
| **EM CD4 Tc** | CD161- | Ly6G- | CD317- | CD5+ | CD3+ | TCRd- | CD4+ | CD44+ | CD62L- |
| **CM/Naive CD4 Tc** | CD161- | Ly6G- | CD317- | CD5+ | CD3+ | TCRd- | CD4+ | CD44- | CD62L+ |
| **CD8 Tc** | CD161- | Ly6G- | CD317- | CD5+ | CD3+ | TCRd- | CD8+ | | |
| **Naive CD8 Tc** | CD161- | Ly6G- | CD317- | CD5+ | CD3+ | TCRd- | CD8+ | CD44+ | CD62L- |
| **CM CD8 Tc** | CD161- | Ly6G- | CD317- | CD5+ | CD3+ | TCRd- | CD8+ | CD44- | CD62L+ |
| **EM CD8 Tc** | CD161- | Ly6G- | CD317- | CD5+ | CD3+ | TCRd- | CD8+ | CD44+ | CD62L+ |

#### Spleen | Peripheral Blood

| Population | Marker Definition | | | | | | | | | | | | | |
|---|---|---|---|---|---|---|---|---|---|---|---|---|---|---|
| **Neutrophils** | CD5- | CD11b+ | Ly6G+ | | | | | | | | | | | |
| **Eosinophils** | CD5- | Ly6G- | SigF+ | SSChi | | | | | | | | | | |
| **Monocytes** | CD5- | Ly6G- | SigF- | SSClo | CD11b+ | Ly6C+ | | | | | | | | |
| **Macrophages** | CD5- | Ly6G- | SigF- | SSClo | CD11b+ | Ly6C- | MHCII+/- | F4/80+ | CD64+ | CD16/32+ | | | | |
| **pDC** | CD5- | Ly6G- | SigF- | SSClo | CD11b+ | Ly6C- | MHCII- | F4/80- | CD317+ | | | | | |
| **Stage I pDC** | CD5- | Ly6G- | SigF- | SSClo | CD11b+ | Ly6C- | MHCII- | F4/80- | CD317+ | CD4- | CD8- | | | |
| **Stage II pDC** | CD5- | Ly6G- | SigF- | SSClo | CD11b+ | Ly6C- | MHCII- | F4/80- | CD317+ | CD4+ | CD8- | | | |
| **Stage III pDC** | CD5- | Ly6G- | SigF- | SSClo | CD11b+ | Ly6C- | MHCII- | F4/80- | CD317+ | CD4+ | CD8+ | | | |
| **cDC** | CD5- | Ly6G- | SigF- | SSClo | CD11b+ | Ly6C- | MHCII+ | F4/80- | CD317- | CD11c+ | | | | |
| **im cDC** | CD5- | Ly6G- | SigF- | SSClo | CD11b+ | Ly6C- | MHCII+ | F4/80- | CD317- | CD11c+ | CCR2- | CD117+ | | |
| **Xcr1 Type im cDC** | CD5- | Ly6G- | SigF- | SSClo | CD11b+ | Ly6C- | MHCII+ | F4/80- | CD317- | CD11c+ | CCR2- | CD117+ | Xcr1+ | CD11b- |
| **CD11b Type im cDC** | CD5- | Ly6G- | SigF- | SSClo | CD11b+ | Ly6C- | MHCII+ | F4/80- | CD317- | CD11c+ | CCR2- | CD117+ | Xcr1- | CD11b+ |
| **mat cDC** | CD5- | Ly6G- | SigF- | SSClo | CD11b+ | Ly6C- | MHCII+ | F4/80- | CD317- | CD11c+ | CCR2- | CD117- | | |
| **Xcr1 Type mat cDC** | CD5- | Ly6G- | SigF- | SSClo | CD11b+ | Ly6C- | MHCII+ | F4/80- | CD317- | CD11c+ | CCR2- | CD117- | Xcr1+ | CD11b- |
| **CD11b+ Type mat cDC** | CD5- | Ly6G- | SigF- | SSClo | CD11b+ | Ly6C- | MHCII+ | F4/80- | CD317- | CD11c+ | CCR2- | CD117- | Xcr1- | CD11b+ |

### Supplemental Table 1 (cont.)

#### Thymus

| Population | Marker Definition | | | | | | | | | |
|---|---|---|---|---|---|---|---|---|---|---|
| **NKc** | CD161+ | CD3e- | | | | | | | | |
| **NK-Tc** | CD161+ | CD3eint | | | | | | | | |
| **DP cells** | CD161- | CD4+ | CD8a+ | | | | | | | |
| **CD4 SP** | CD161- | CD4+ | CD8a- | | | | | | | |
| **CD8 SP** | CD161- | CD4- | CD8a+ | | | | | | | |
| **DN** | CD161- | CD4- | CD8a- | | | | | | | |
| **TN** | CD161- | CD4- | CD8a- | CD3e- | gd- | | | | | |
| **TN1** | CD161- | CD4- | CD8a- | CD3e- | gd- | CD44+ | CD25- | | | |
| **ETP (TN1a/b)** | CD161- | CD4- | CD8a- | CD3e- | gd- | CD44+ | CD25- | CD24+ | CD117hi | |
| **TN1c** | CD161- | CD4- | CD8a- | CD3e- | gd- | CD44+ | CD25- | CD24+ | CD117int | |
| **TN1d** | CD161- | CD4- | CD8a- | CD3e- | gd- | CD44+ | CD25- | CD24+ | CD117- | |
| **TN1e** | CD161- | CD4- | CD8a- | CD3e- | gd- | CD44+ | CD25- | CD24- | CD117- | |
| **TN2** | CD161- | CD4- | CD8a- | CD3e- | gd- | CD44+ | CD25+ | CD24+ | CD117+ | |
| **TN3a** | CD161- | CD4- | CD8a- | CD3e- | gd- | CD44- | CD25+ | CD24+ | CD71- | |
| **TN3b** | CD161- | CD4- | CD8a- | CD3e- | gd- | CD44- | CD25+ | CD24+ | CD71+ | |
| **TN4** | CD161- | CD4- | CD8a- | CD3e- | gd- | CD44- | CD25- | CD24+ | | |
| **immature DP** | CD161- | CD4+ | CD8a+ | CD3e- | gd- | CD44- | CD25- | CD24+ | CD71+ | |
| **small resting DP** | CD161- | CD4+ | CD8a+ | CD3e- | gd- | CD44- | CD25- | CD24+ | CD71- | |
| **CD3- CD69+ DP** | CD161- | CD4+ | CD8a+ | CD3e- | gd- | CD44- | CD25- | CD24+ | CD71- | CD69+ |
| **CD3+ CD69+ DP** | CD161- | CD4+ | CD8a+ | CD3e+ | gd- | CD44- | CD25- | CD24+ | CD71- | CD69+ |
| **CD3+ CD4 SP** | CD161- | CD4+ | CD8a- | CD3e+ | gd- | CD44- | | | | |
| **Treg** | CD161- | CD4+ | CD8a- | CD3e- | gd- | CD44- | CD25+ | | | |
| **Early CD4 SP** | CD161- | CD4+ | CD8a- | CD3e- | gd- | CD44- | CD25+ | CD24hi | | |
| **Late CD4 SP** | CD161- | CD4+ | CD8a- | CD3e- | gd- | CD44- | CD25+ | CD24lo | QA-1+ | |
| **CD3+ CD8 SP** | CD161- | CD4- | CD8a+ | CD3e+ | gd- | CD44- | CD25- | CD24+ | | |
| **Early CD8 SP** | CD161- | CD4- | CD8a+ | CD3e+ | gd- | CD44- | CD25- | CD24+ | | |
| **Late CD8 SP** | CD161- | CD4- | CD8a+ | CD3e+ | gd- | CD44- | CD25- | CD24lo | QA-1+ | |
| **gd-Tc** | CD161- | CD4- | CD8a+ | CD3e+ | gd+ | | | | | |
| **DN3a gd-Tc** | CD161- | CD4- | CD8a+ | CD3e+ | gd+ | CD25+ | CD71+ | | | |
| **DN4a gd-Tc** | CD161- | CD4- | CD8a+ | CD3e+ | gd+ | CD25- | CD44- | CD71+ | | |
| **DN4b gd-Tc** | CD161- | CD4- | CD8a+ | CD3e+ | gd+ | CD25- | CD44- | | | |
| **Cluster A gd Tc** | CD161- | CD4- | CD8a+ | CD3e+ | gd+ | CD44+ | CD24- | | | |
| **Cluster B gd Tc** | CD161- | CD4- | CD8a+ | CD3e+ | gd+ | CD44- | CD24+ | | | |

#### Peritoneal Cavity

| Populations | Marker definition | | | | | | |
|---|---|---|---|---|---|---|---|
| **Bc** | CD19+ | | | | | | |
| **B1a** | CD19+ | CD5+ | CD11b+/- | | | | |
| **B1b** | CD161+ | CD5- | CD11b+ | | | | |
| **B2** | CD161+ | CD5- | CD11b- | | | | |
| **Neutrophils** | CD19- | CD5- | CD11b+ | Ly6G+ | | | |
| **Mast Cells** | CD19- | CD5- | CD11b- | CD117+ | | | |
| **Eosinophils** | CD19- | CD5- | CD11b+ | CD117- | SigF+ | SSChi | Ly6C+/- |
| **Macrophages** | CD19- | CD5- | CD11b- | CD117- | SigF- | SSChi | F4/80+ |
| **Tc** | CD19- | CD5+ | MHCII- | | | | |
| **Ly6C+ Tc** | CD19- | CD5+ | MHCII- | Ly6C+ | | | |
| **NKc** | CD19- | CD161+ | CD5- | | | | |
| **Ly6C+ NKc** | CD19- | CD161+ | CD5- | Ly6C+ | | | |
| **imDC** | CD19- | CD11c+ | MHCII+ | CD117+ | | | |
| **Myelomonocyte** | CD19- | CD11b+ | CD117- | | | | |
| **Myelomonocyte P1** | CD19- | CD11b+ | CD117- | CD192+ | Ly6C+ | | |
| **Myelomonocyte P2** | CD19- | CD11b+ | CD117- | CD192+ | Ly6C+ | MHCII+ | |
| **Myelomonocyte P3** | CD19- | CD11b+ | CD117- | CD192+ | Ly6C- | MHCII+ | |

**Supplemental Table 2.** Panel of antibody clones for immunophenotyping

| | Specificity | Clone | Provider |
|---|---|---|---|
| Thymus | CD003e | 2C11 | eBioscience |
| | CD004 | RM4.5 | BD Biosciences |
| | CD005 | 53-7.3 | BD Biosciences |
| | CD008a | 53-6.7 | BD Biosciences |
| | CD024 | M1.69 | BD Biosciences |
| | CD025 | PC61 | Biolegend |
| | CD027 | LG.3A10 | BD Biosciences |
| | CD044 | IM7 | Biolegend |
| | CD069 | H1.2F3 | BD Biosciences |
| | CD071 | R17217 | Biolegend |
| | CD117 | 2B8 | BD Biosciences |
| | CD161 | PK136 | BD Biosciences |
| | TCRd | GL3 | eBioscience |
| | QA-2 | 695H1.9.9 | Biolegend |
| Spleen | CD003e | 2C11 | BD Biosciences |
| | CD004 | RM4.5 | BD Biosciences |
| | CD005 | 53-7.3 | BD Biosciences |
| | CD008 | 53-6.7 | BD Biosciences |
| | CD011b | M1.70 | BD Biosciences |
| | CD011c | HL3 | BD Biosciences |
| | CD019 | 1D3 | BD Biosciences |
| | CD044 | IM7 | BD Biosciences |
| | CD062L | MEL14 | Biolegend |
| | CD161 | PK136 | BD Biosciences |
| | CD317 | 927 | eBioscience |
| | Ly6C | AL21 | BD Biosciences |
| | Ly6G | 1A8 | BD Biosciences |
| | TCRd | GL3 | eBioscience |
| | F4/80 | BM8 | Biolegend |
| | MHCII | M5-114.15.2 | Biolegend |
| | IgD | 11-26c.2A | Biolegend |
| Peritoneum | CD011c | HL3 | BD Biosciences |
| | LY6g | 1A8 | BD Biosciences |
| | CD005 | 53-7.3 | BD Biosciences |
| | MHCII | M5/114.15.2 | Biolegend |
| | CD117 | 2B8 | BD Biosciences |
| | CD161 | PK136 | BD Biosciences |
| | CD011b | M1/70 | BD Biosciences |
| | LY6c | AL-21 | BD Biosciences |
| | CD115 | AFS98 | ThermoFisher Sc |
| | CD192 | SA203G11 | Biolegend |
| | SigF | E50-2440 | BD Biosciences |
| | CD19 | 1D3 | ThermoFisher Sc |
| | F4/80 | BM8 | Biolegend |
| | CD064 | X54-5/7.1 | Biolegend |
| PBL | CD005 | 53-7.3 | BD Biosciences |
| | CD008a | 53-6.7 | BD Biosciences |
| | CD019 | 6D5 | Biolegend |
| | CD011b | M1/70 | BD Biosciences |
| | CD044 | IM7 | BD Biosciences |
| | CD045 | 30F11 | ThermoFisher Sc |
| | CD004 | RM4-5 | BD Biosciences |
| | CD161 | PK136 | BD Biosciences |
| | Ly6G | 1A8 | BD Biosciences |
| | Ly6C | HK1.4 | Biolegend |
